# Protective antigen-mediated delivery of an anti-CRISPR protein for precision genome editing

**DOI:** 10.1101/2025.02.23.639680

**Authors:** Axel O. Vera, Nicholas L. Truex, Vedagopuram Sreekanth, Bradley L. Pentelute, Amit Choudhary, Ronald T. Raines

## Abstract

Precise control over the dosage of Cas9-based technologies is essential because off-target effects, mosaicism, chromosomal aberrations, immunogenicity, and genotoxicity can arise with prolonged Cas9 activity. Type II anti-CRISPR proteins (Acrs) inhibit and control Cas9 but are generally impermeable to the cell membrane due to their size (6–54 kDa) and anionic charge. Moreover, existing Acr delivery methods are long-lived and operate within hours (*e.g*., viral and non-viral vectors) or are not applicable in vivo (*e.g*., nucleofection), limiting therapeutic applications. To address these problems, we developed the first protein-based anti-CRISPR delivery platform, LF_N_-Acr/PA, which delivers Acrs into cells within minutes. LF_N_-Acr/PA is a nontoxic, two-component protein system derived from anthrax toxin, where protective antigen proteins bind receptors widespread in human cells, forming a pH-triggered endosomal pore that LF_N_-Acr binds and uses to enter the cell. In the presence of PA, LF_N_-Acr enters human cells at concentrations as low as 2.5 pM to inhibit up to 95% of Cas9-mediated knockout, knock-in, transcriptional activation, and base editing. Timing LF_N_-Acr delivery reduces off-target base editing and increases Cas9 specificity by 41%. LF_N_-Acr/PA is the most potent known cell-permeable CRISPR-Cas inhibition system, significantly improving the utility of CRISPR for genome editing.

## Introduction

Clustered regularly interspaced short palindromic repeats (CRISPR)/CRISPR-associated (Cas) systems use RNA-guided endonucleases (*e.g*., Cas9) to cleave DNA and defend bacteria against phages (1, 2). Cas9 has been repurposed as a genome-editing technology in human cells because of its programmability (3, 4). Cas9-based base editors (5, 6) and prime editors (7) have also been developed to modify DNA sequences without double-strand breaks. Cas9 technologies are used to treat genetic disorders (8-10), but their use is limited because they can cause unintended DNA modifications (*i.e*., off-target effects) (11-14), genetic mosaicism (15), chromosomal aberrations (16, 17), immunogenicity (18), and genotoxicity (19, 20).

Reducing the half-life of Cas9 in cells can overcome these limitations. We (21-24) and others (25) have developed small-molecule inhibitors (*e.g*., BRD7586), degraders, and activators that control the half-life of Cas9 and increase its specificity, defined as the on-target to off-target ratio of DNA modification. Because of their size (<800 Da) and lipophilicity, small-molecule Cas9 modulators are cell-permeable and rapidly act on Cas9. These small molecules, however, exhibit low aqueous solubility, interact weakly with Cas9, and bind and degrade other proteins indiscriminately (25, 26). Furthermore, most modulators interact only with Cas9 variants fused to small-molecule-binding domains. These domains reduce activity and production yield of Cas9, complicate its cellular delivery, and are incompatible with Cas9–effector protein fusions used for precision genome editing.

Unlike small molecules, type II anti-CRISPR proteins (Acrs) bind Cas9 strongly and selectively because phages evolved them to prevent bacterial CRISPR immunity (27, 28). Acrs have been shown to inhibit Cas9 and Cas9–effector protein fusions by preventing any of the three stages of Cas9 interference: gRNA loading, DNA binding, or DNA cleavage (29). For example, AcrIIA4, an anionic Acr, binds the PAM interacting domain and RuvC nuclease domain of the Cas9-gRNA ribonucleoprotein (RNP) complex to prevent DNA binding (30-32). Acrs have been used to control Cas9 (33, 34), mitigate its toxicity (35), and increase its efficiency (36) and specificity (32, 37-41).

Because of their cellular impermeability, Acrs require delivery vehicles to enter cells. DNA encoding Acrs has been delivered by using viral vectors (*e.g*., adenovirus (35), AAVs (37, 38, 42), and lentiviruses (43, 44)), lipid transfection (27, 28), and nucleofection (32, 40). Acr mRNA transcripts have been delivered via microinjection (45) and lipid nanoparticles (46). The AAV and mRNA delivery methods confirmed that Acrs are functional in vivo and have therapeutic potential (38, 45, 46). AAVs are, however, immunogenic and lead to the persistent production of the Acr via episome formation in the nucleus or host genome integration (47). In turn, mRNA delivery provides transient activity and limits the risk of host genome integration but still requires translation of the transcript, which requires time and can be impacted by cell stress and type. The slow delivery kinetics of DNA and mRNA challenge the control of Acr activity and delay Cas9 inhibition to hours beyond administration (48). In contrast, an Acr delivered as a protein could inhibit Cas9 immediately.

Nucleofection has been used to deliver Acrs in the protein form into the cytosol and nucleus (32). As a non-viral strategy, nucleofection avoids host genome integration, prolonged production of the Acr, and packaging limitations. Nucleofection does require an external chamber and electrical pulses and is limited to ex vivo applications. Further, the sequential delivery of Cas9 and Acrs using nucleofection could introduce other sources of stress and toxicity to cells (49).

Here, we describe the first protein-based delivery system for introducing Acrs into human cells. We show that an Acr fused to the nontoxic N-terminal domain of lethal factor from *Bacillus anthracis* (LF_N_-Acr) enters cells in the presence of protective antigen (PA)—a pore-forming protein. LF_N_-Acr/PA is the most potent known cell-permeable CRISPR-Cas inhibition system, combining the exquisite target specificity of an Acr with the cell permeability of small molecules to inhibit Cas9-based technologies efficiently and increase genome-editing specificity.

## Results

### Design of the Cellular Delivery System

PA is an 83-kDa pore-forming protein used by the anthrax lethal toxin from the Gram-positive bacterium, *B. anthracis*, to deliver the cytotoxic protein lethal factor (LF) into the cytosol of mammalian cells via the following mechanism: First, PA binds to the anthrax receptors, ANTXR1 and ANTXR2 (50, 51), which are displayed on the surface of most human cells (52). Second, proteolytic cleavage by a cell-surface furin protease activates PA by converting it into a 63 kDa protein (PA_63_) (53). Third, PA_63_ oligomerizes into a heptameric or octameric prepore on the cell surface (54, 55). Fourth, LF binds to the prepore with nanomolar affinity (56, 57). Fifth, acidification of the endosome triggers pore formation and translocation of LF into the cytosol (58-60). We hypothesized that AcrIIA4 covalently fused to the nontoxic N-terminal domain of lethal factor (LF_N_), LF_N_-Acr, would enter cells in the presence of PA (Fig. 1). We and others have used PA to deliver different proteins (52), including apo-Cas9 (61). The LF_N_/PA delivery platform has many advantages, including being compatible in vivo (62-64), non-viral, short-lived (65, 66), effective at <20-min delivery timescales (67), and optionally bioreversible (66, 68) and tissue-specific (64, 69-71).

**Fig. 1.**
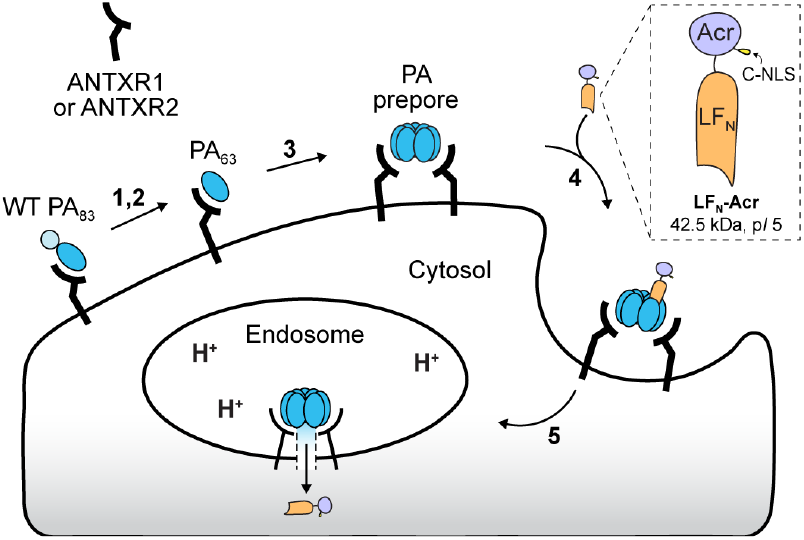
Mechanism of the Protective antigen (PA)-mediated delivery of LF_N_-Acr into cells. (1) Wild-type PA (*i.e*., PA 83 kDa: PA_83_) binds to the ANTXR1 or ANTXR2 receptors on the cell surface. (2) Cell-surface proteases cleave and activate receptor-bound PA_83_. (3) The activated PA (*i.e*., PA 63 kDa: PA_63_) oligomerizes into a prepore. (4) The nontoxic N-terminal domain of lethal factor (LF_N_) in the LF_N_-Acr fusion binds to the PA prepore. (5) The entire complex is endocytosed. Acidification of the endosome promotes the PA prepore to change into a transmembrane pore, which translocates LF_N_-Acr into the cytosol.

We produced and purified LF_N_-Acr using an MBP-TEV system in *Escherichia coli* and assessed purity via SDS–PAGE and Q-TOF LC–MS (Fig. S1). LF_N_-Acr is a 42.5-kDa anionic protein (p*I* 5, Fig. 1). For reference, wild-type AcrIIA4 is a 10.2-kDa anionic protein (p*I* 4). We chose AcrIIA4 as a model Acr because it potently inhibits *Streptococcus pyogenes* Cas9 (SpCas9) and tolerates modifications at both of its termini (31).

### LF_N_-Acr Enters Cells in the Presence of PA and Inhibits Cas9 Nuclease Activity in a Dose- and Time-Dependent Manner

We tested LF_N_-Acr delivery by monitoring Cas9 nuclease activity via the *GFP*-disruption assay and benchmarked its efficiency against BRD7586, the most potent known small-molecule Cas9 inhibitor (23). We delivered Cas9 RNP, targeting a *GFP* gene, via nucleofection into a cell line stably producing a GFP–PEST fusion (U2OS-EGFP.PEST) (72), incubated the cells for 48 h, and imaged. Throughout the incubation (*e.g*., at a 2-h dosing time), we added LF_N_-Acr and PA or BRD7586 (Fig. 2*A*). Cas9-mediated cleavage of *GFP* reduces GFP fluorescence. Conversely, inhibition of Cas9 with LF_N_-Acr conserves GFP fluorescence. LF_N_-Acr (2.5 pM–2.5 µM) in the presence of PA (20 nM) inhibited Cas9 activity (always normalized to 100% without Acr) from 100–8.7% (Figs. 2*B, C* and S2–S6). LF_N_-Acr (2.5 µM) alone did not inhibit Cas9, confirming that PA mediates the delivery of LF_N_-Acr. BRD7586 (2.5–20 µM) reduced Cas9 activity from 100–81% (Figs. 2*B*, S7, and S8).

**Fig. 2.**
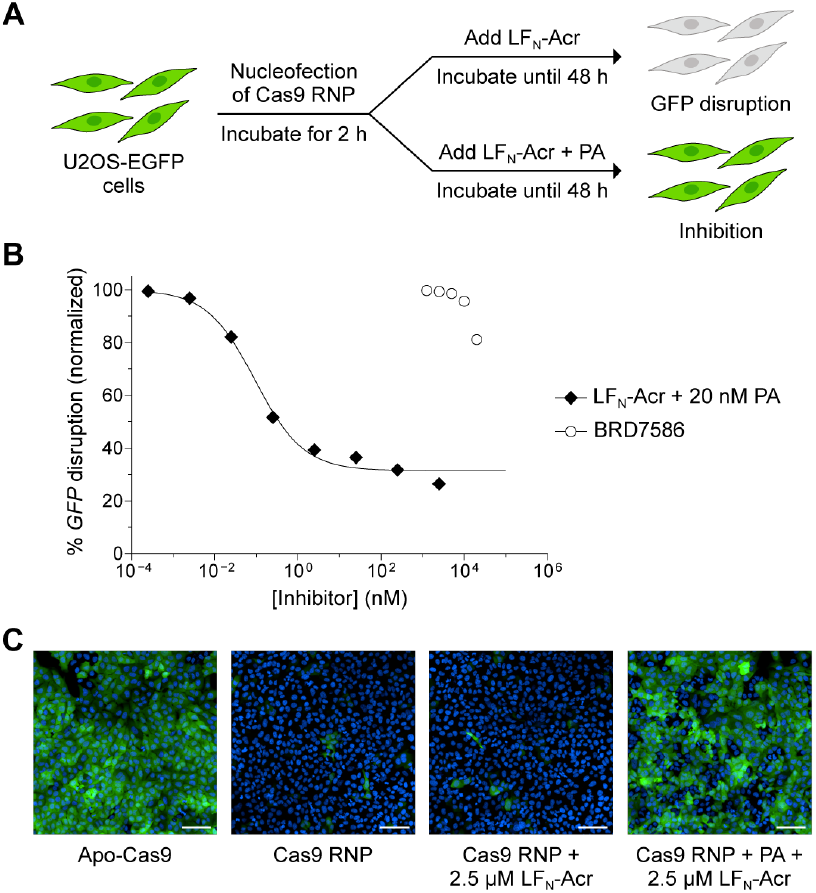
Effect of LF_N_-Acr delivery on *GFP* disruption. (*A*) A cartoon showing the experimental set-up for the *GFP*-disruption assay. The PA-mediated delivery of LF_N_-Acr prevents *GFP* disruption. (*B*) Dose-dependent inhibition of Cas9 by LF_N_-Acr and BRD7586 in the *GFP*-disruption assay. C-NLS-SpCas9 RNP (20 pmol), targeting *GFP*, was delivered via nucleofection into U2OS-EGFP.PEST cells. The cells were incubated at 37 °C and dosed with LF_N_-Acr (2.5 × 10^−4^, 2.5 × 10^−3^, 0.025, 0.25, 2.5, 25, 250, and 2,500 nM) and PA (20 nM) or BRD7586 (1.25, 2.5, 5, 10, and 20 µM) at a 2-h dosing time. After 48 h, live cells were stained with Hoechst 33342 and imaged via high-throughput confocal microscopy. The values were normalized to Cas9 RNP (100%, not shown) and are the mean ± standard deviation (SD) of three independent replicates. The error bars are too small to visualize. (*C*) Representative fluorescence images from the *GFP*-disruption assay. Controls include nucleofection of Cas9 RNP followed by dosing with LF_N_-Acr (2.5 µM) at a 2-h dosing time. The cell nucleus was stained with Hoechst 33342 (blue). Scale bars, 100 µm.

LF_N_-Acr in the presence of PA (IC_5_ = 0.0031 nM, Table S1) inhibits Cas9 3.2 × 10^6^-fold more potently than BRD7586 (IC_5_ = 10,000 nM, Table S1) when comparing the doses that inhibit Cas9 activity by 5% (IC_5_). We used IC_5_ to compare LF_N_-Acr and BRD7586 activity because BRD7586 is cytotoxic at concentrations >20 µM. Because the value of IC_50_ is necessarily greater than that of IC_5_, LF_N_-Acr (IC_50_ = 0.35 nM, Table S1) inhibits Cas9 >2.9 × 10^4^-fold more potently than BRD7586 in our assay conditions. Thus, LF_N_-Acr/PA is the most potent cell-permeable CRISPR-Cas inhibition system.

We monitored Cas9 nuclease activity via the *HiBiT*-knock-in assay to validate LF_N_-Acr delivery further. We delivered Cas9 RNP, targeting *GAPDH*, and a *HiBiT* single-stranded oligodeoxynucleotide (ssODN) donor via nucleofection into HEK293T cells, incubated the cells for 72 h while adding LF_N_-Acr and PA at dosing times ranging from 0–48 h, and quantified luminescence (Fig. 3*A*). Cas9-mediated cleavage of *GAPDH* and insertion of the *HiBiT* ssODN via homology-directed repair (HDR) leads to the formation of NanoLuc in the presence of LgBiT. Conversely, inhibition of Cas9 with LF_N_-Acr prevents NanoLuc formation. LF_N_-Acr (250 pM– 2.5 µM) in the presence of PA (20 nM) reduced Cas9 activity from 100–17% (Figs. 3*B, C*, S9, and S10). LF_N_-Acr (2.5 µM) alone did not inhibit Cas9. We also assessed cell viability during the HiBiT assay to confirm that the PA-mediated delivery of LF_N_-Acr is not toxic (Fig. S11).

**Fig. 3.**
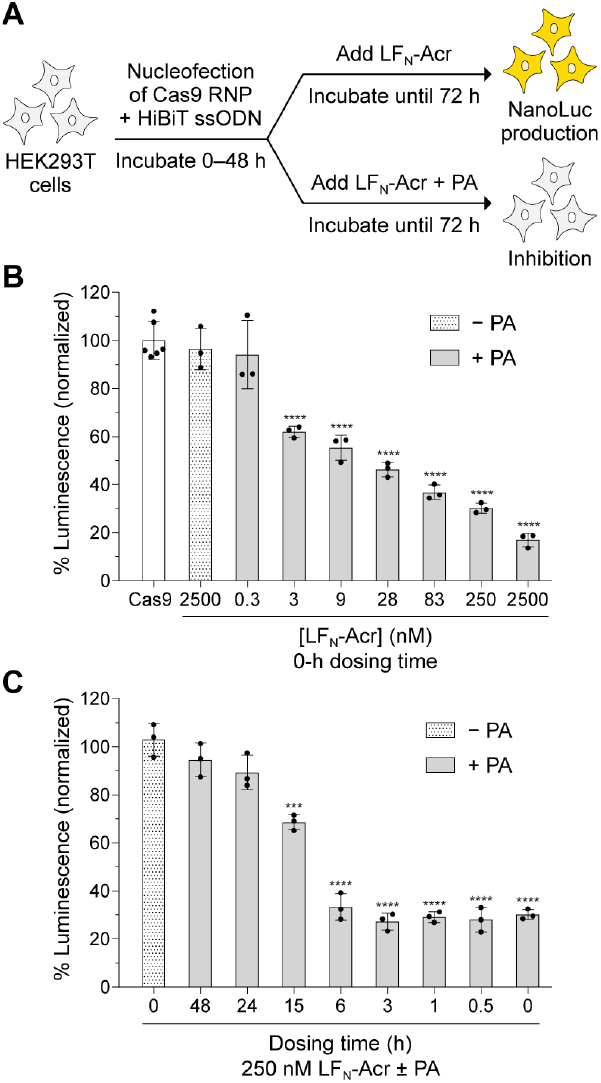
Effect of LF_N_-Acr delivery on *HiBiT* knock-in. (*A*) A cartoon showing the experimental set-up for the *HiBiT*-knock-in assay. The PA-mediated delivery of LF_N_-Acr prevents Cas9-mediated knock-in and NanoLuc formation. (*B, C*) Dose- and time-dependent inhibition of Cas9 by LF_N_-Acr in the *HiBiT-*knock-in assay. C-NLS-SpCas9 RNP (20 pmol), targeting *GAPDH*, and the *HiBiT* ssODN (80 pmol) were co-delivered via nucleofection into HEK293T cells. The cells were incubated at 37 °C and dosed with LF_N_-Acr (0.3–2,500 nM) and PA (20 nM) at a 0-h dosing time (*B*) or dosed with LF_N_-Acr (250 nM) and PA (20 nM), varying the dosing time from 0–48 h (*C*). After 72 h, the cells were lysed and combined with LgBiT, and their luminescence was quantified. Controls include co-nucleofection of Cas9 RNP and ssODN (white bar, panel *B*) and co-nucleofection of Cas9 RNP and ssODN followed by dosing with 2,500 nM LF_N_-Acr (− PA, panel *B*) or 250 nM LF_N_-Acr (− PA, panel *C*) at a 0-h dosing time. The values were normalized to Cas9 RNP + ssODN and are the mean ± standard deviation (SD) of three independent replicates. The significance of LF_N_-Acr/PA additions was determined with an unpaired, two-tailed *t*-test versus Cas9 RNP + ssODN, where *, **, ***, and **** refer to *P* ≤ 0.05, *P* ≤ 0.01, *P* ≤ 0.001, and *P* ≤ 0.0001, respectively.

LF_N_-Acr is less potent in the *HiBiT* assay (IC_50_ = 0.71–17 nM, Table S1) than in the *GFP* assay (IC_50_ = 0.35–2.9 nM, Table S1). This difference is expected because the HEK293T cells used in the *HiBiT* assay display fewer copies of the PA-targeted receptors than the U2OS cells used in the *GFP* assay (73-75). The *HiBiT* assay detects inhibition less precisely than the *GFP* assay at lower LF_N_-Acr concentrations, further contributing to the apparent decrease in potency.

### LF_N_-Acr Inhibits CRISPR-Mediated Transcriptional Activation (CRISPRa) by dCas9-VPR

Recombinant Acr production has been used to inhibit dCas9-based transcriptional regulation in cells (28, 33, 44), suggesting that LF_N_-Acr would also control this process. Hence, we used nucleofection to deliver a dCas9-VPR plasmid into HEK293T cells stably transcribing sgRNAs complementary to a CRE-promoter upstream of NanoLuc. We then incubated the cells for 24 h while adding LF_N_-Acr and PA at dosing times ranging from 0–12 h and quantified luminescence (Fig. 4*A*). dCas9-VPR binding of the CRE promoter induces transcription of NanoLuc and increases luminescence. Inhibition of dCas9-VPR with LF_N_-Acr decreases luminescence. LF_N_-Acr (25 nM–2.5 µM) in the presence of PA (20–100 nM) reduced dCas9-VPR activity from 100–5.5% (Figs. 4*B* and S12). LF_N_-Acr (2.5 µM) alone did not inhibit dCas9-VPR.

**Fig. 4.**
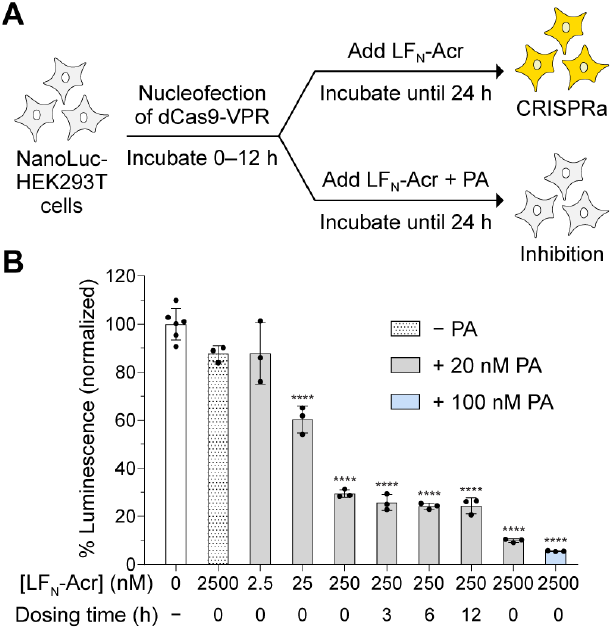
Effect of LF_N_-Acr delivery on CRISPRa. (*A*) A cartoon showing the experimental set-up for the CRISPRa assay. The PA-mediated delivery of LF_N_-Acr prevents dCas9-VPR-mediated production of NanoLuc. (*B*) Dose- and time-dependent inhibition of dCas9-VPR by LF_N_-Acr in the CRISPRa assay. dSpCas9-VPR plasmid (300 ng) was delivered via nucleofection into 7×sgRNA-CRE-NanoLuc-HEK293T cells. The cells were incubated at 37 °C and dosed with LF_N_-Acr (2.5– 2,500 nM) and PA (20–100 nM) after 0–12 h (dosing time). After 24 h, the cells were incubated with furimazine, and their luminescence was quantified. Controls include nucleofection of dCas9-VPR plasmid (white bar) and nucleofection of dCas9-VPR plasmid followed by dosing with 2,500 nM LF_N_-Acr (− PA) at a 0-h dosing time. The values were normalized to dCas9-VPR plasmid and are the mean ± SD of three independent replicates. The significance of LF_N_-Acr/PA additions was determined with an unpaired, two-tailed *t*-test versus dCas9-VPR, where *, **, ***, and **** refer to *P* ≤ 0.05, *P* ≤ 0.01, *P* ≤ 0.001, and *P* ≤ 0.0001, respectively.

LF_N_-Acr inhibited dCas9-VPR in a dose-dependent manner at a constant PA concentration (Fig. 4*B*). At a constant LF_N_-Acr concentration, increasing the PA concentration increased dCas9-VPR inhibition (Fig. 4*B*). Increasing the concentration of PA may be beneficial in applications where a vector constantly produces Cas9 or when the cells display fewer copies of the PA-targeted receptors.

LF_N_-Acr inhibited dCas9-VPR with equal potency at a 0-h and 12-h dosing time (Fig. 4*B*), suggesting that cells require at least 12 h to produce dCas9-VPR from a plasmid. Furthermore, inhibition by dCas9-VPR at a 12-h dosing time allows for the displacement of dCas9-VPR from the CRE promoter (*e.g*., by cellular factors) and its subsequent inhibition by LF_N_-Acr. The reversibility of CRISPRa and fast degradation of NanoLuc-PEST (half-life: 20 min) (76) could also explain why dCas9-VPR inhibition remains potent with time and suggests that LF_N_-Acr remains active in cells for at least 24 h.

### LF_N_-Acr Increases Cas9 Specificity

Nucleofection of AcrIIA4 6 h after introducing Cas9 increases Cas9 specificity in cells (32), suggesting that varying the dosing time of LF_N_-Acr would have a similar effect. To verify this hypothesis, we delivered Cas9 RNP, targeting *EMX1*, via nucleofection into HEK293T cells, incubated the cells for 72 h while adding LF_N_-Acr and PA at dosing times ranging from 0–48 h, and used next-generation sequencing (NGS) to assess Cas9 cleavage at the *EMX1* on-target and off-target sites (Fig. 5*A*). LF_N_-Acr (3–250 nM) in the presence of PA (20 nM) reduced Cas9 on-target activity from 100–60% and off-target activity from 42–18% (Fig. S13). LF_N_-Acr (250 nM) alone did not inhibit Cas9. The inhibition of Cas9 by LF_N_-Acr increased Cas9 specificity by 1.11-to 1.41-fold or 11–41% relative to uninhibited Cas9 (Fig. 5*B, C* and Eqs. S1–S3). We also assessed the Cas9-mediated cleavage of *EMX1* via the T7 endonuclease 1 (T7E1) mismatch detection assay (Fig. S14). This experiment further validates that PA mediates LF_N_-Acr delivery in a dose- and time-dependent manner (Fig. S15).

**Fig. 5.**
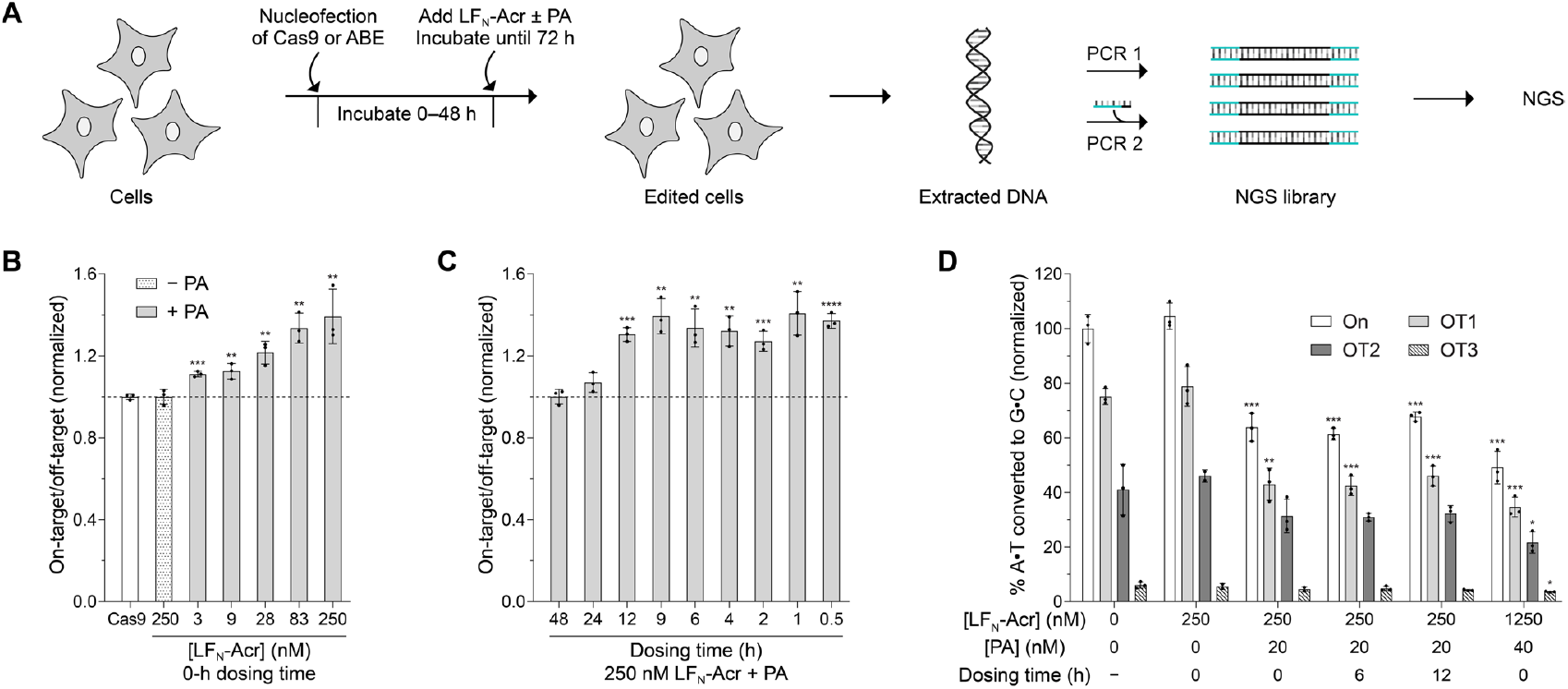
Effect of LF_N_-Acr delivery on genome-editing specificity. (*A*) A cartoon showing the experimental set-up for LF_N_-Acr delivery experiments involving next-generation sequencing (NGS). (*B, C*) LF_N_-Acr delivery increases Cas9 specificity in a dose- and time-dependent manner. NLS-SpCas9-NLS RNP (80 pmol), targeting *EMX1*, was delivered via nucleofection into HEK293T cells. The cells were incubated at 37 °C and dosed with LF_N_-Acr (3–250 nM) and PA (20 nM) at a 0-h dosing time (*B*) or dosed with LF_N_-Acr (250 nM) and PA (20 nM), varying the dosing time from 0.5–48 h (*C*). After 72 h, genomic DNA was extracted, sequenced via NGS, and analyzed using the CRISPResso2 software pipeline to determine insertions, deletions, and substitutions (% modification) at the on-target and off-target sites. Controls include nucleofection of Cas9 RNP (white bar, panel *B*) and nucleofection of Cas9 RNP followed by dosing with 250 nM LF_N_-Acr (− PA, panel *B*) at a 0-h dosing time. The specificity was calculated as the on-target to off-target % modification ratio, normalizing to the on-target to off-target % modification ratio of Cas9 RNP. The significance of the specificity was determined with an unpaired, two-tailed *t*-test versus the specificity of Cas9 RNP, where *, **, ***, and **** refer to *P* ≤ 0.05, *P* ≤ 0.01, *P* ≤ 0.001, and *P* ≤ 0.0001, respectively. (*D*) LF_N_-Acr delivery regulates base editing. ABE8e (500 ng) and sgRNA (165 ng) plasmids, targeting *HBG2*, were delivered via nucleofection into HEK293T cells. The cells were incubated at 37 °C and dosed with LF_N_-Acr (250–1250 nM) and PA (20–40 nM) after 0–12 h (dosing time). After 72 h, genomic DNA was extracted, sequenced via NGS, and analyzed using CRISPResso2 to determine the % modification at the on-target (On) and off-target (OT1, OT2, and OT3) sites. Controls include nucleofection of ABE8e-sgRNA plasmids (− dosing time) and nucleofection of ABE8e-sgRNA plasmids followed by dosing with LF_N_-Acr (250 nM) at a 0-h dosing time. The values were normalized to ABE8e-sgRNA (On) and are the mean ± SD of three independent replicates. The significance of LF_N_-Acr/PA additions was determined with an unpaired, two-tailed *t*-test versus ABE8e-sgRNA (On, OT1, OT2, and OT3), where *, **, ***, and **** refer to *P* ≤ 0.05, *P* ≤ 0.01, *P* ≤ 0.001, and *P* ≤ 0.0001, respectively.

### LF_N_-Acr Regulates Base Editing

Acr plasmids delivered via nucleofection inhibit base editors (40). To verify if LF_N_-Acr also inhibits base editing, we delivered ABE8e and sgRNA plasmids, targeting *HBG2*, via nucleofection into HEK293T cells, incubated the cells for 72 h while adding LF_N_-Acr and PA at dosing times ranging from 0–12 h, and used NGS to assess on-target and off-target A to G conversions. LF_N_-Acr (250–1,250 nM) in the presence of PA (20–40 nM) reduced ABE8e on-target activity from 100–49% and off-target activity from 75–35, 41–35, and 6.1–3.6% at off-target sites 1, 2, and 3, respectively (Fig. 5*D*). LF_N_-Acr (250 nM) alone did not inhibit ABE8e. LF_N_-Acr reduced guide-dependent off-target base editing. Similar to Acr plasmids (40) and small-molecule Cas9 degraders (22, 24), LF_N_-Acr did not increase base editing specificity significantly (Fig. S16).

## Discussion

This work introduces means to use Acrs to inhibit Cas9-based technologies and increase genome-editing specificity in human cells. The advantages of our Acr delivery method include speed, efficiency, and short-lived exposure. PA mediates LF_N_-Acr delivery even at picomolar concentrations. Engineered PA variants can also be used to target specific cell receptors and cell types (62, 69-71). For example, we recently conjugated PA to a single-chain variable fragment and a full-length immunoglobulin G antibody to target dendritic and cancer cells, respectively (64, 77). We could deliver LF_N_-Acr to specific cells using one of these PA variants. Cell-specific Acr delivery applications might be beneficial when Cas9 exhibits off-tissue editing, e.g., to reduce the risk of unintended germline editing in the ovaries while editing hepatocytes in the liver in vivo (9).

LF_N_-Acr can be used to probe Cas9 kinetics. For example, our NGS studies show that the PA-mediated delivery of LF_N_-Acr at a 24-h dosing time inhibits Cas9 RNP (Fig. S13), suggesting that Cas9 RNP remains active in cells for at least 24 h. In addition, LF_N_-Acr delivery inhibits plasmid-encoded ABE8e-sgRNA similarly at a 0-h and 12-h dosing time (Fig. 5*D*), suggesting that cells require at least 12 h to produce ABE8e-sgRNA from plasmids.

Timing the delivery of LF_N_-Acr increases Cas9 specificity by up to 41% (Fig. 5*B, C*). Due to the limitations of target-amplicon sequencing via NGS, we only assessed cleavage at a known *EMX1* off-target site. Therefore, 41% is an underrepresentation of the Cas9 specificity increase obtained with LF_N_-Acr. We expect genome-wide Cas9 off-target analyses (e.g., GUIDE-Seq (78)) to uncover the full potential of LF_N_-Acr in the future.

The LF_N_-Acr/PA delivery system enables us to use dose and time to optimize genome-editing specificity at different targets. Our system can also be further optimized for precision genome editing. For example, we could engineer LF_N_-Acr variants with milder activity or different mechanisms of action to inhibit Cas9 off-target activity with higher selectivity, as shown using Acr vectors (39, 79, 80).

LF_N_-Acr did not significantly increase the specificity of ABE8e (Fig. S16), probably because the kinetics of ABE8e guide-dependent on-target and off-target base editing are similar in the *HBG2* site. Our small-molecule-mediated ABE8e degradation studies at the *HBG2* site support this hypothesis (22, 24). Nonetheless, LF_N_-Acr is likely to increase the specificity of base editors incorporating kinetically slower deaminases, as shown with deaminase inhibitor proteins (81). LF_N_-Acr could also be used to reduce bystander editing, that is, base editing proximal to the targeted nucleotide, as has been done with Acr peptides in vivo (82).

In summary, we established the first protein-based delivery system for introducing Acrs into human cells. The LF_N_-Acr/PA delivery system combines the target specificity of an Acr with the cell permeability of small molecules to allow rapid and on-demand Cas9 control. We showed that LF_N_-Acr inhibits Cas9 nuclease, dCas9-VPR-based CRISPRa systems, and base editors efficiently and increases Cas9 specificity. Ongoing work focuses on determining the efficiency and immunogenicity of the LF_N_-Acr/PA delivery system in vivo. We aim to validate its use as a co-therapy for therapeutic genome editing applications. The LF_N_-Acr/PA delivery system could be used to reduce off-target and off-tissue genome editing in vivo and enhance genome editing therapies in the future.

## Materials and Methods

### Conditions

Unless indicated otherwise, all procedures were performed in air at ambient temperature (∼22 °C) and pressure (1.0 atm).

### Recombinant DNA

DNA fragments encoding the sequences of LF_N_ and Cys-AcrIIA4-NLS were inserted downstream of an N-terminal 10×His tag, maltose binding protein (MBP), and a TEV protease cleavage site in a custom vector (Addgene plasmid #115670) (83) via Gibson assembly to generate the LF_N_-Acr bacterial production and purification plasmid (pAV14). LF_N_-Acr contains an engineered cysteine residue to enable site-specific chemical conjugation in future applications.

### Protein Purification

Protein purification proceeded as described previously (83, 84) with modifications. The LF_N_-Acr plasmid (pAV14) was grown in *E. coli* Rosetta2 (DE3) cells in Lysogeny Broth supplemented with 100 µg/mL ampicillin and 2% w/v glucose at 37 °C for 16 h. The starter culture was subcultured in Terrific Broth supplemented with 100 µg/mL ampicillin and 2% w/v glucose at 37 °C until the OD_600_ was 0.6–0.8. Protein production was induced with 500 μM isopropyl-β-D-thiogalactoside (IPTG), and the cultures were grown at 16 °C for 16 h. The cells were harvested, resuspended in lysis buffer (20 mM Tris–HCl buffer, pH 7.5, containing 500 mM NaCl, 20 mM imidazole, 1 mM tris(2-carboxyethyl)phosphine (TCEP), and 5% v/v glycerol) supplemented with 0.5 mM phenylmethanesulfonyl fluoride and Roche cOmplete Protease Inhibitor Cocktail, and lysed by sonication. MBP–LF_N_-Acr was purified using Ni-NTA Superflow resin (Qiagen) in lysis buffer supplemented with either 20 mM imidazole (wash) or 300 mM imidazole (elution). Eluted MBP–LF_N_-Acr was cleaved with TEV protease (purified in-house using Addgene plasmid #8827 (85)) overnight at 4 °C and dialyzed into Acr buffer (20 mM Tris– HCl buffer, pH 7.5, containing 150 mM NaCl, 1 mM TCEP, and 5% v/v glycerol). Cleaved MBP– LF_N_-Acr was loaded onto an MBPTrap HP (Cytiva) upstream of a HiTrap Q HP (Cytiva) and eluted over a linear gradient of NaCl (0.150–1.00 M) in 20 mM Tris–HCl buffer, pH 7.5, containing 1 mM TCEP and 5% v/v glycerol. LF_N_-Acr was then loaded onto a HiLoad 26/600 Superdex 200 pg (Cytiva) and eluted with Acr buffer. Purified LF_N_-Acr was concentrated to 250 µM in Acr buffer and snap-frozen in N_2_(l) for storage at −80 °C. LF_N_-Acr concentration was measured using *A*_280 nm_ and a BCA assay kit (Thermo Scientific). LF_N_-Acr purity was assessed by SDS–PAGE (Coomassie blue staining) and Q-TOF LC–MS.

### Cell Culture

U2OS cells with an integrated *GFP*–*PEST* fusion gene (U2OS-EFFP.PEST) and HEK293T cells were cultured in cell culture medium (Dulbecco’s modified Eagle’s medium with GlutaMax (DMEM, Gibco) supplemented with 10% v/v fetal bovine serum (FBS, Corning), 100 U/mL penicillin−streptomycin (Gibco), and 1 mM sodium pyruvate (Gibco)) at 37 °C in a 5% v/v CO_2_(g) atmosphere. HEK293T cells stably expressing 7 different sgRNAs complementary to CRE-promoter regions upstream of NanoLuc (7×sgRNA-CRE-NanoLuc-HEK293T) were cultured in cell culture medium supplemented with puromycin dihydrochloride (2 µg/mL, Gibco). Puromycin was removed before performing experiments. All Acr delivery experiments were performed in cell culture medium containing FBS (10% v/v).

### Acr Addition/Delivery to Cells

A 0-h dosing time is defined as the time when LF_N_-Acr and PA are added to Cas9-nucleofected cells before starting the incubation at 37 °C. A >0-h dosing time implies adding LF_N_-Acr and PA to Cas9-nucleofected cells >0 h after starting the incubation at 37 °C. For dose-dependency studies at a 2-h dosing time, LF_N_-Acr (250 fM–2.5 µM) and PA (20 nM) were added to the cells 2 h after Cas9 delivery by replacing the cell culture medium in the well with fresh medium containing LF_N_-Acr and PA diluted to the desired concentration. The Acr buffer in the medium was <1% v/v at all doses in this medium-exchange method. For dose-dependency studies at a 0-h dosing time or for time-dependency studies at a constant concentration, LF_N_-Acr (250 fM–2.5 µM) and PA (20–100 nM) were added to the cells 0–48 h after Cas9 delivery by directly pipetting a concentrated LF_N_-Acr/PA solution in Acr buffer to the cell culture medium in the well. The concentration of the LF_N_-Acr/PA solution in Acr buffer in this direct addition method was adjusted so that the final concentration of Acr buffer in the medium was 4.8% v/v (*e.g*., 5 µL of an LF_N_-Acr/PA solution added to 100 µL of cell culture medium). BRD7586 (1.25–20 µM, MedChemExpress), dissolved in DMSO, was added to cells using the medium-exchange and direct addition methods described previously, keeping the DMSO concentration constant at 0.4% v/v.

### *GFP*-Disruption Assay

SpCas9-C-NLS RNP (20 pmol), targeting the *GFP* gene, was delivered via nucleofection into U2OS-EGFP.PEST cells (300,000 cells) using an SE Cell Line 4D-Nucleofector X Kit (Lonza, DN-100 protocol). The nucleofected cells were resuspended in cell culture medium (100,000 cells/mL), transferred to a 96-well plate (100 µL), and incubated at 37 °C. 0–36 h after starting the incubation (dosing time), LF_N_-Acr (250 fM–2.5 µM) and PA (20 nM) were added to the cells via the medium-exchange (2-h dosing time) or direct addition method (all other dosing times). After 48 h, live cells were stained with 2 µg/mL Hoechst 33342 (37 °C, 20 min) in FluoroBrite DMEM (Gibco) and imaged via high throughput confocal microscopy (Perkin Elmer Opera Phenix; 488 and 561 nm excitation for fluorescence, 740 nm transmitted light for digital phase contrast). Cas9-mediated cleavage disrupts the *GFP* gene. Successful delivery of LF_N_-Acr prevents Cas9-mediated *GFP* disruption.

### HiBiT Knock-In Assay

SpCas9-C-NLS RNP (20 pmol), targeting the *GAPDH* gene, and a single-stranded oligo donor nucleotide (ssODN) encoding the HiBiT tag (80 pmol) were delivered via nucleofection into HEK293T cells (300,000 cells) using an SF Cell Line 4D-Nucleofector X Kit (Lonza, CM-130 protocol). The nucleofected cells were resuspended in cell culture medium (150,000 cells/mL), transferred to a poly(D-lysine)-coated 96-well plate (100 µL), and incubated at 37 °C. 0–48 h after starting the incubation (dosing time), LF_N_-Acr (2.5 pM–2.5 µM) and PA (20 nM) were added to the cells via the medium-exchange (2-h dosing time) or direct addition method (all other dosing times). After 72 h, the cells were incubated with the PrestoBlue HS reagent, and their viability was measured via fluorescence (560 nm excitation/590 nm emission, SpectraMax M5). The cells were lysed in the presence of LgBiT and lytic substrate (Nano-Glo HiBiT Lytic Detection System, Promega), and their luminescence was quantified (Perkin Elmer EnVison). Cas9-mediated cleavage and HiBiT tag insertion via homology-directed repair produce NanoLuc after LgBiT addition. Successful delivery of LF_N_-Acr prevents Cas9-mediated cleavage, HiBiT tag insertion, and NanoLuc formation.

### CRISPRa Assay

A plasmid encoding dSpCas9-VPR (Addgene plasmid #63798 (86)) was delivered via nucleofection into 7×sgRNA-CRE-NanoLuc-HEK293T cells (300,000 cells) using an SF Cell Line 4D-Nucleofector X Kit (Lonza, CM-130 protocol). The nucleofected cells were resuspended in cell culture medium (350,000 cells/mL), transferred to a poly(D-lysine)-coated 96-well plate (100 µL), and incubated at 37 °C. 0–12 h after starting the incubation (dosing time), LF_N_-Acr (25 pM–2.5 µM) and PA (20–100 nM) were added to the cells via direct addition of a concentrated LF_N_-Acr/PA solution (5 µL). After 24 h, the cells were incubated with the PrestoBlue HS reagent, and their viability was measured via fluorescence (560 nm excitation/590 nm emission, SpectraMax M5). Then, the cells were incubated with 10 µM furimazine (37 °C, 15 min), and their luminescence was quantified (Perkin Elmer EnVison). The binding of dCas9-VPR to the CRE promoter produces NanoLuc and luminescence. Successful delivery of LF_N_-Acr prevents dCas9-VPR from binding to the CRE promoter and NanoLuc formation.

### Next-Generation Sequencing (Cas9 RNP)

NLS-SpCas9-NLS RNP (80 pmol), targeting the *EMX1* gene, was delivered via nucleofection into HEK293T cells (300,000 cells) using an SF Cell Line 4D-Nucleofector X Kit (Lonza, CM-130 protocol). The nucleofected cells were resuspended in cell culture medium (240,000 cells/mL), transferred to a 24-well poly(D-lysine)-coated plate (571.4 µL), and incubated at 37 °C. 0–48 h after starting the incubation (dosing time), LF_N_-Acr (3– 250 nM) and PA (20 nM) were added to the cells via direct addition of a concentrated LF_N_-Acr/PA solution (28.6 µL). After 72 h, genomic DNA was extracted (QIAamp DNA Mini Kit), and the target and off-target sites were PCR amplified, barcoded, and purified via gel extraction (MinElute Gel Extraction Kit). The amplicons were sequenced via NGS and analyzed using the CRISPResso2 software pipeline to determine insertions, deletions, and substitutions (% modification).

### T7 Endonuclease 1 (T7E1) Assay

C-NLS-SpCas9 RNP (20 pmol), targeting the *EMX1* gene, was delivered via nucleofection into HEK293T cells (300,000 cells) using an SF Cell Line 4D-Nucleofector X Kit (Lonza, CM-130 protocol). The nucleofected cells were resuspended in cell culture medium (240,000 cells/mL), transferred to a 24-well poly(D-lysine)-coated plate (571.4 µL), and incubated at 37 °C. 0–24 h after starting the incubation (dosing time), LF_N_-Acr (30 pM–250 nM) and PA (20 nM) were added to the cells via direct addition of a concentrated LF_N_-Acr/PA solution (28.6 µL). After 72 h, genomic DNA was extracted (QIAamp DNA Mini Kit), and the target site was PCR amplified, rehybridized, incubated with T7E1 (NEB), and visualized via agarose gel electrophoresis.

### Next-Generation Sequencing (ABE8e plasmid)

ABE8e (500 ng, Addgene plasmid #138489) and sgRNA (165 ng) plasmids, targeting the *HBG2* gene (87), were delivered via nucleofection into HEK293T cells (300,000 cells) using an SF Cell Line 4D-Nucleofector X Kit (Lonza, CM-130 protocol). The nucleofected cells were resuspended in cell culture medium (230,000 cells/mL), transferred to a 24-well poly(D-lysine)-coated plate (571.4 µL), and incubated at 37 °C. 0–12 h after starting the incubation (dosing time), LF_N_-Acr (250–1,250 nM) and PA (20–40 nM) were added to the cells via direct addition of a concentrated LF_N_-Acr/PA solution (28.6 µL). After 72 h, genomic DNA was extracted (QIAamp DNA Mini Kit), and the target and off-target sites were PCR amplified, barcoded, and purified via gel extraction (MinElute Gel Extraction Kit). The amplicons were sequenced via NGS and analyzed using the CRISPResso2 software pipeline to determine insertions, deletions, and substitutions (% modification).

## Acknowledgments

The authors thank Benjamin K. Law (Broad Institute of MIT and Harvard) for providing NLS-SpCas9-NLS protein and Prof. J. Keith Joung (Harvard Medical School) for providing the U2OS-EGFP.PEST cell line. They are grateful to Drs. Maria Alimova and Aditya Raguram (Broad Institute of MIT and Harvard) for providing training in imaging and next-generation sequencing, respectively. A.O.V. was supported by an MIT Dean of Science Fellowship, an NIH predoctoral fellowship (F31 GM148042), and an HHMI Gilliam fellowship (GT15945). N.L.T. was supported by an NIH postdoctoral fellowship (F32 CA239362). Cell imaging was performed with an Opera Phenix High-Throughput imaging system that was supported by Grant S10 OD026839 (NIH). This work was supported by Grants R01 GM137606 to A.C. and R.T.R. and R35 GM148220 (NIH) to R.T.R.

