## Supplementary Material for "Protective antigen-mediated delivery of an anti-CRISPR protein for precision genome editing"

**Supplementary Figures**

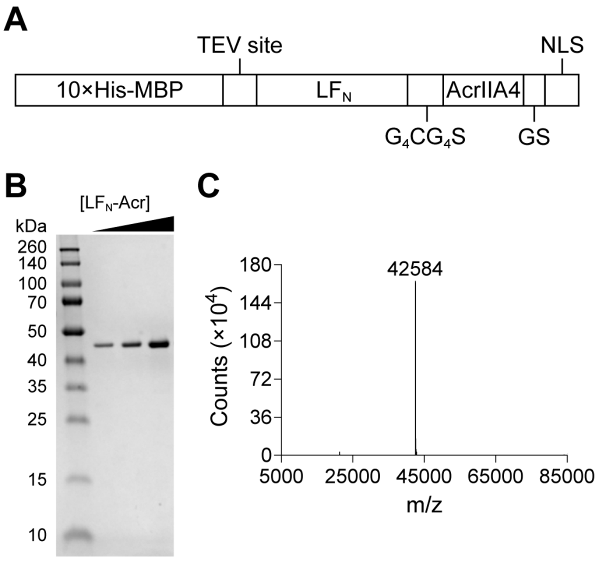


**Fig. S1.** Purification of $LF_N$-Acr. (*A*) The construct that was used to produce and purify $LF_N$-Acr. A protein fusion between a polyhistidine-tagged maltose-binding protein (MBP, for solubility and purification), a TEV protease cleavage site, a glycine–cysteine–serine linker ($G_4CG_4S$, for flexibility and optional bioconjugation), the N-terminus of lethal factor ($LF_N$, for binding to PA), AcrIIA4 (for Cas9 inhibition), a glycine–serine (GS) linker (for flexibility), and an SV40 nuclear localization signal (NLS). (*B*) Assessing $LF_N$-Acr purity via sodium dodecyl sulfate–polyacrylamide gel electrophoresis (SDS–PAGE). The concentration of $LF_N$-Acr increases from left to right. (*C*) Assessing $LF_N$-Acr purity via quadrupole time-of-flight liquid-chromatography–mass spectrometry (Q-TOF LC–MS).

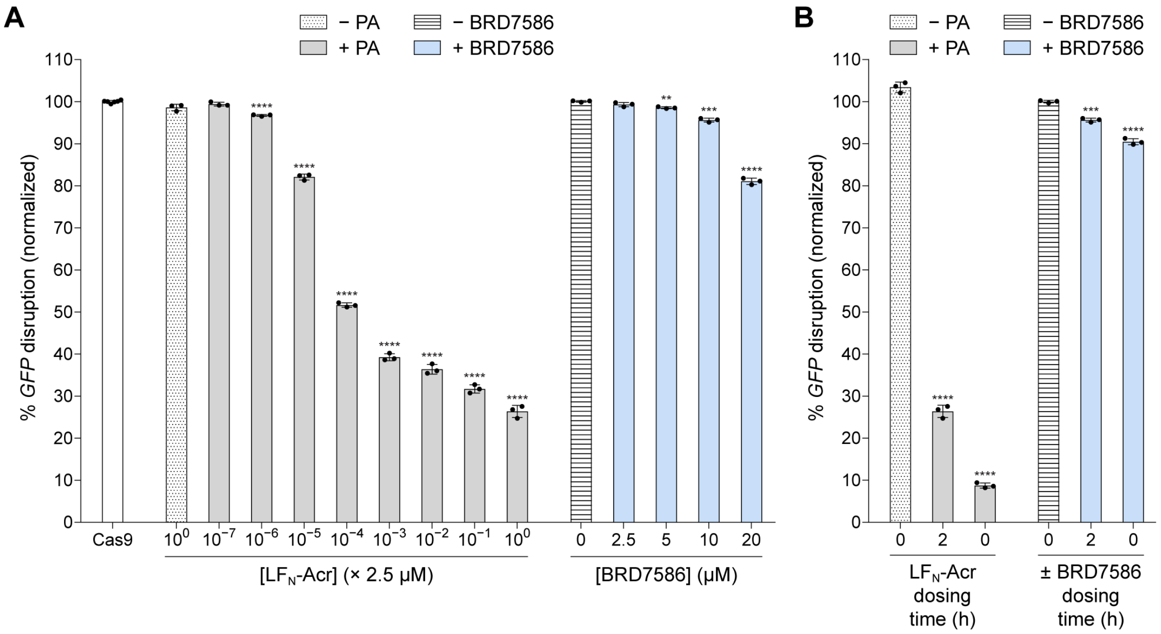


**Fig. S2.** Dose- and time-dependent inhibition of Cas9 by $LF_N$-Acr and BRD7586 in the *GFP*-disruption assay. C-NLS-SpCas9 RNP (20 pmol), targeting *GFP*, was delivered via nucleofection into U2OS-EGFP.PEST cells. The cells were incubated at 37 °C and dosed with $LF_N$-Acr (2.5 × $10^{-7}$–2.5 μM) and PA (20 nM) or BRD7586 (2.5–20 μM) at a 2-h dosing time (*A*) or dosed with $LF_N$-Acr (2.5 μM) and PA (20 nM) or BRD7586 (10 μM), varying the dosing time from 0–2 h (*B*). After 48 h, live cells were stained with Hoechst 33342 and imaged via high throughput confocal microscopy. Controls include nucleofection of Cas9 RNP (white bar) and nucleofection of Cas9 RNP followed by dosing with 2.5 μM $LF_N$-Acr (− PA) or 0.4% v/v DMSO (− BRD7586) at a 2-h dosing time (panel *A*) or dosing with 2.5 μM $LF_N$-Acr (− PA) or 0.4% v/v DMSO (− BRD7586) at a 0-h dosing time (panel *B*). The values were normalized to Cas9 RNP ($LF_N$-Acr) or DMSO (BRD7586) and are the mean ± standard deviation (SD) of three independent replicates. The significance of $LF_N$-Acr/PA or BRD7586 additions was determined with an unpaired, two-tailed *t*-test versus Cas9 RNP ($LF_N$-Acr/PA) or DMSO (BRD7586), where *, **, ***, and **** refer to $P \leq 0.05$, $P \leq 0.01$, $P \leq 0.001$, and $P \leq 0.0001$, respectively.

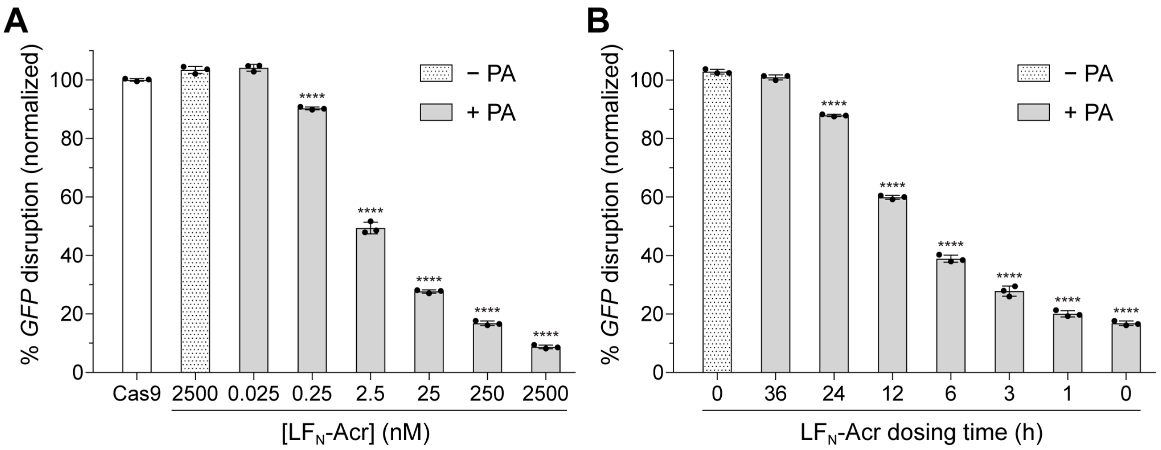


**Fig. S3.** Dose- and time-dependent inhibition of Cas9 by $LF_N$-Acr in the *GFP*-disruption assay. C-NLS-SpCas9 RNP (20 pmol), targeting *GFP*, was delivered via nucleofection into U2OS-EGFP.PEST cells. The cells were incubated at 37 °C and dosed with $LF_N$-Acr (0.025–2,500 nM) and PA (20 nM) at a 0-h dosing time (*A*) or dosed with $LF_N$-Acr (250 nM) and PA (20 nM), varying the dosing time from 0–36 h (*B*). After 48 h, live cells were stained with Hoechst 33342 and imaged via high throughput confocal microscopy. Controls include nucleofection of Cas9 RNP (white bar) and nucleofection of Cas9 RNP followed by dosing with 2,500 nM $LF_N$-Acr (− PA, panel *A*) or 250 nM $LF_N$-Acr (− PA, panel *B*) at a 0-h dosing time. The values were normalized to Cas9 RNP and are the mean ± SD of three independent replicates. The significance of $LF_N$-Acr/PA additions was determined with an unpaired, two-tailed *t*-test versus Cas9 RNP, where *, **, ***, and **** refer to $P \leq 0.05$, $P \leq 0.01$, $P \leq 0.001$, and $P \leq 0.0001$, respectively.

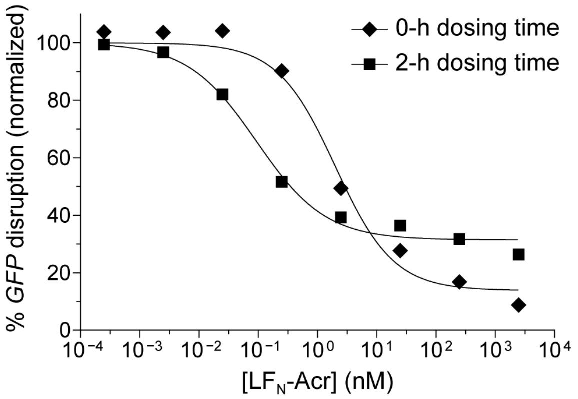


**Fig. S4.** Effect of dosing time in the dose-dependent inhibition of Cas9 by $LF_N$-Acr in the *GFP*-disruption assay. C-NLS-SpCas9 RNP (20 pmol), targeting *GFP*, was delivered via nucleofection into U2OS-EGFP.PEST cells. The cells were incubated at 37 °C and dosed with $LF_N$-Acr ($2.5 \times 10^{-4}$, $2.5 \times 10^{-3}$, 0.025, 0.25, 2.5, 25, 250, and 2,500 nM) and PA (20 nM) at a 0- or 2-h dosing time. After 48 h, live cells were stained with Hoechst 33342 and imaged via high throughput confocal microscopy. The values were normalized to Cas9 RNP (100%, not shown) and are the mean ± SD of three independent replicates. The error bars are too small to visualize.

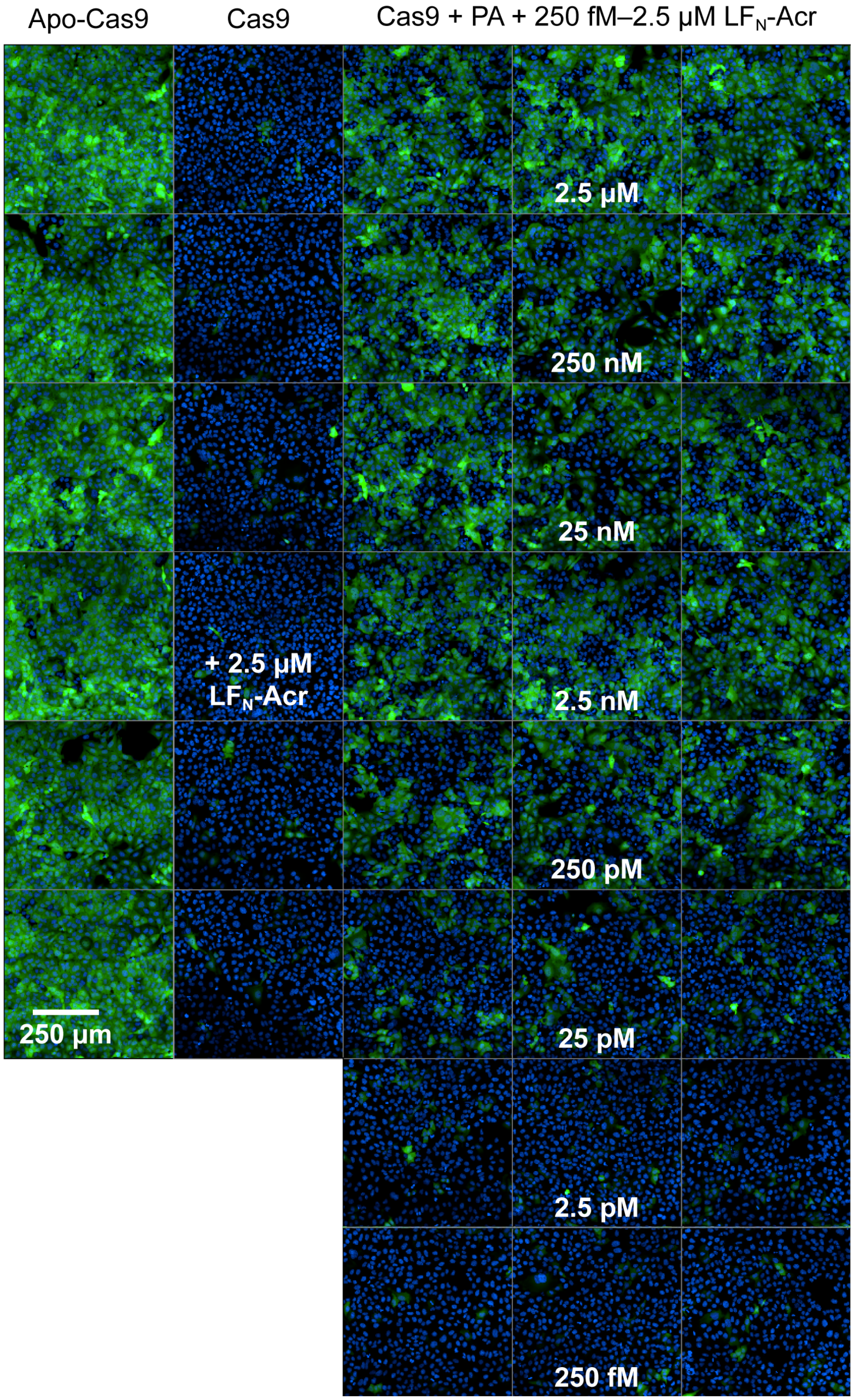


**Fig. S5.** Representative fluorescence images from the *GFP*-disruption assay in Fig. S2*A* show the dose-dependent inhibition of Cas9 by LF$_N$-Acr (250 fM–2.5 μM) in the presence of PA (20 nM) at a 2-h dosing time. Controls include nucleofection of Apo-Cas9, nucleofection of Cas9 RNP, and nucleofection of Cas9 RNP followed by dosing with LF$_N$-Acr (2.5 μM) at a 2-h dosing time. The nucleus appears blue due to staining with Hoechst 33342.

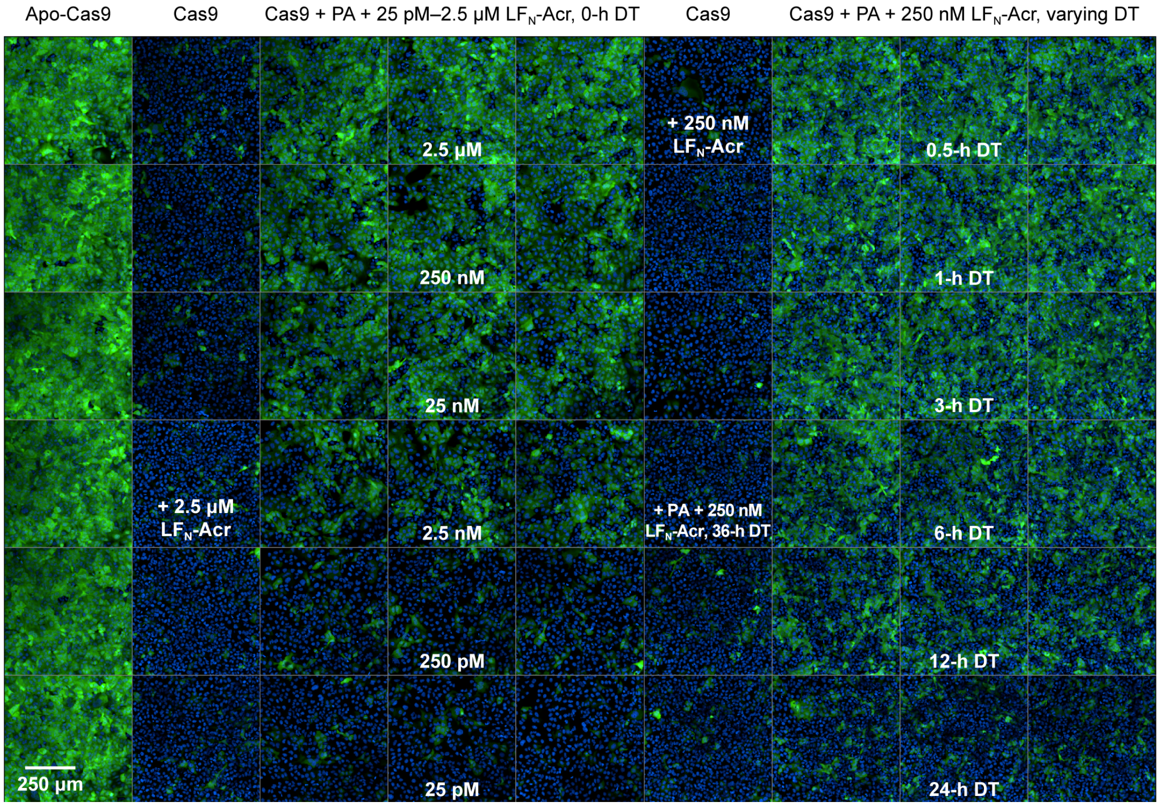


**Fig. S6.** Representative fluorescence images from the *GFP*-disruption assay in Fig. S3 show the dose-dependent inhibition of Cas9 by LF$_N$-Acr (25 pM–2.5 μM) in the presence of PA (20 nM) and time-dependent inhibition of Cas9 by LF$_N$-Acr (250 nM) in the presence of PA (20 nM). Controls include nucleofection of Apo-Cas9, nucleofection of Cas9 RNP, and nucleofection of Cas9 RNP followed by dosing with LF$_N$-Acr (250 nM and 2.5 μM) at a 0-h dosing time (DT). The nucleus appears blue due to staining with Hoechst 33342.

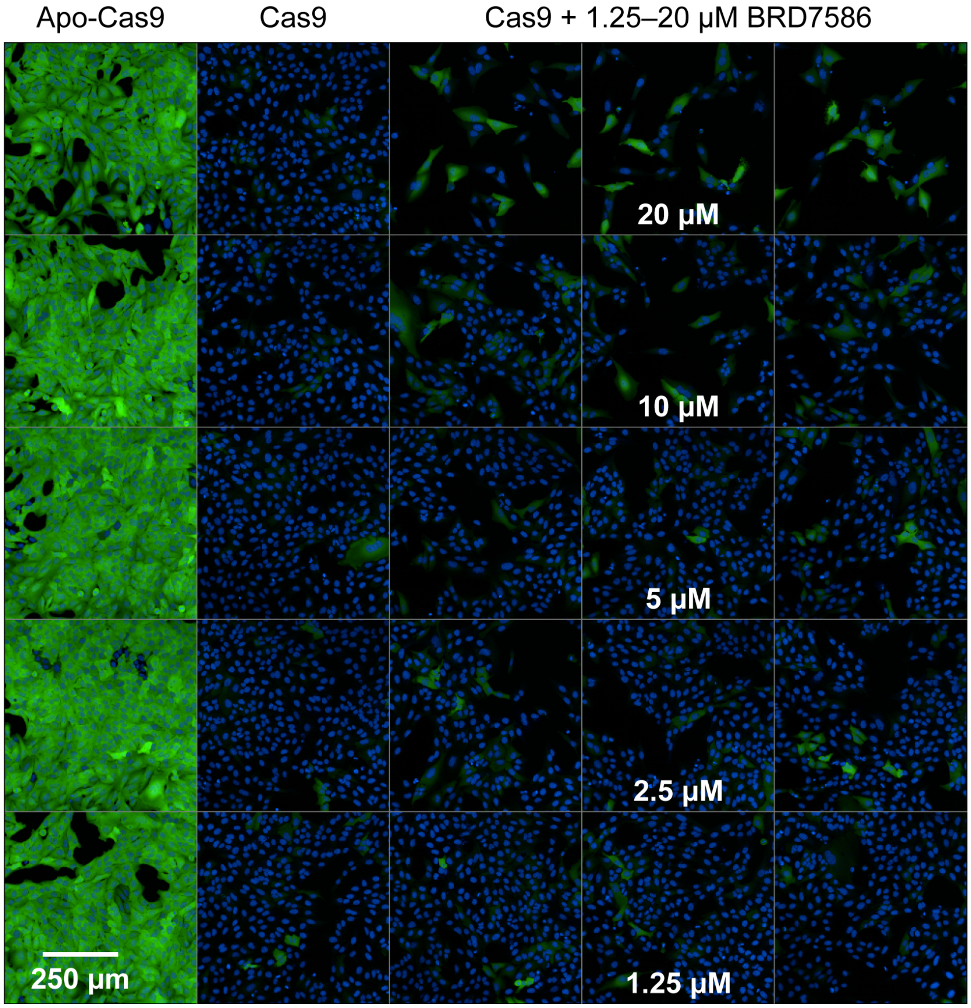


**Fig. S7.** Representative fluorescence images from the *GFP*-disruption assay in Fig. S2*A* show the dose-dependent inhibition of Cas9 by BRD7586 (1.25–20 µM) at a 2-h dosing time. Controls include nucleofection of Apo-Cas9 and nucleofection of Cas9 RNP followed by dosing with DMSO (0.4% v/v) at a 2-h dosing time. The nucleus appears blue due to staining with Hoechst 33342.

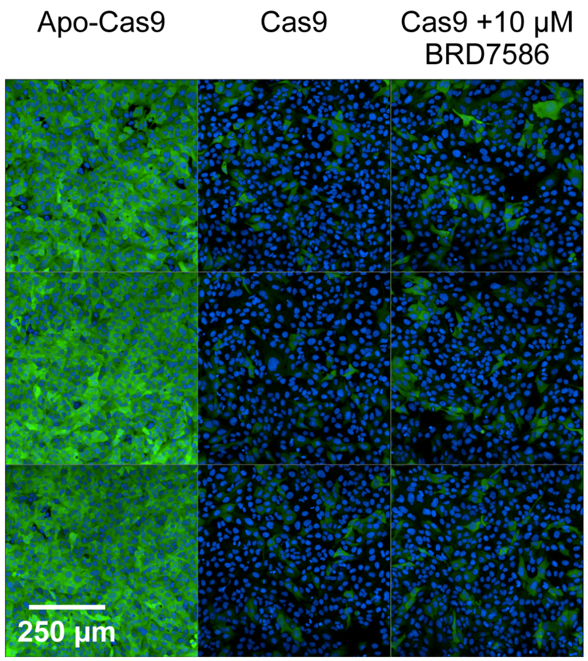


**Fig. S8.** Representative fluorescence images from the *GFP*-disruption assay in Fig. S2*B* show the inhibition of Cas9 by BRD7586 (10 µM) at a 0-h dosing time. Controls include nucleofection of Apo-Cas9 and nucleofection of Cas9 RNP followed by dosing with DMSO (0.4% v/v) at a 0-h dosing time. The nucleus appears blue due to staining with Hoechst 33342.

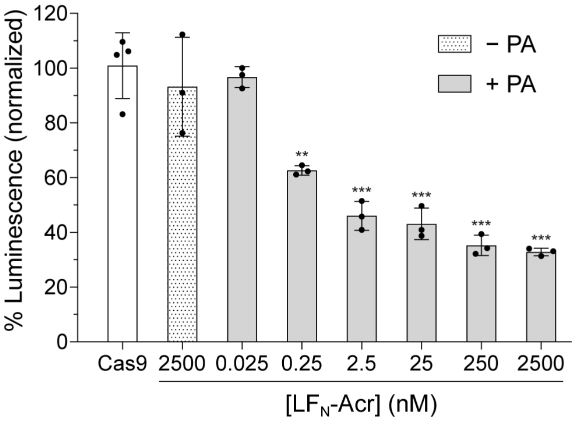


**Fig. S9.** Dose-dependent inhibition of Cas9 by $LF_N$-Acr in the *HiBiT*-knock-in assay. C-NLS-SpCas9 RNP (20 pmol), targeting *GAPDH*, and the *HiBiT* ssODN (80 pmol) were co-delivered via nucleofection into HEK293T cells. The cells were incubated at 37 °C and dosed with $LF_N$-Acr (0.025–2,500 nM) and PA (20 nM) at a 2-h dosing time. After 72 h, the cells were lysed and combined with LgBiT, and their luminescence was quantified. Controls include co-nucleofection of Cas9 RNP and ssODN (white bar) and co-nucleofection of Cas9 RNP and ssODN followed by dosing with 2,500 nM $LF_N$-Acr (− PA) at a 2-h dosing time. The values were normalized to Cas9 RNP + ssODN and are the mean ± SD of three independent replicates. The significance of $LF_N$-Acr/PA additions was determined with an unpaired, two-tailed *t*-test versus Cas9 RNP + ssODN, where *, **, ***, and **** refer to $P \leq 0.05$, $P \leq 0.01$, $P \leq 0.001$, and $P \leq 0.0001$, respectively.

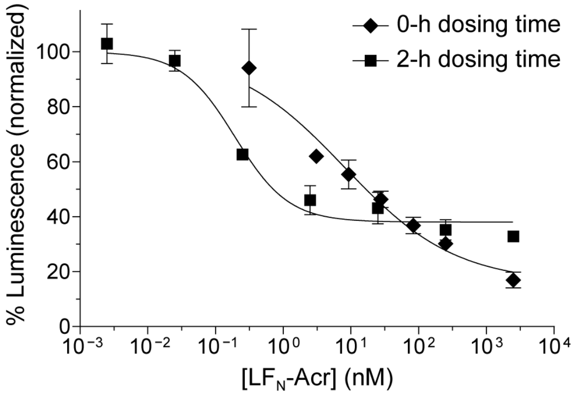


**Fig. S10.** Effect of dosing time in the dose-dependent inhibition of Cas9 by $LF_N$-Acr in the *HiBiT*-knock-in assay. C-NLS-SpCas9 RNP (20 pmol), targeting *GAPDH*, and the *HiBiT* ssODN (80 pmol) were co-delivered via nucleofection into HEK293T cells. The cells were incubated at 37 °C and dosed with PA (20 nM) and $LF_N$-Acr (0.3, 3, 9, 28, 83, 250, and 2,500 nM at a 0-h dosing time or $2.5 \times 10^{-3}$, 0.025, 0.25, 2.5, 25, 250, and 2,500 nM at a 2-h dosing time). After 72 h, the cells were lysed and combined with LgBiT, and their luminescence was quantified. The values were normalized to Cas9 RNP + ssODN (100%, not shown) and are the mean ± SD of three independent replicates. Some error bars are too small to visualize.

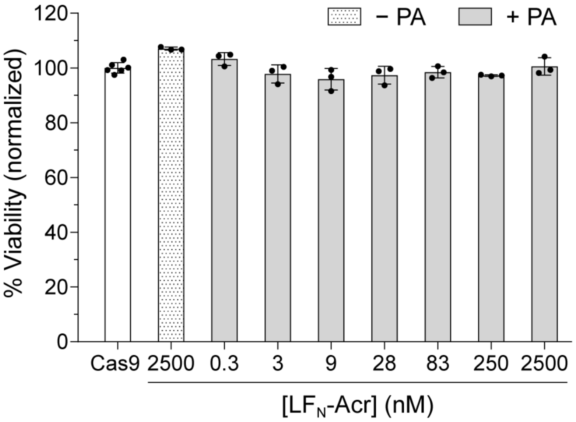


**Fig. S11.** Cell viability after $LF_N$-Acr delivery in the *HiBiT*-knock-in assay. C-NLS-SpCas9 RNP (20 pmol), targeting *GAPDH*, and the *HiBiT* ssODN (80 pmol) were co-delivered via nucleofection into HEK293T cells. The cells were incubated at 37 °C and dosed with $LF_N$-Acr (0.3–2,500 nM) and PA (20 nM) at a 0-h dosing time. After 72 h, the cells were incubated with the PrestoBlue HS reagent. Cell viability was determined by measuring fluorescence (560 nm excitation/590 nm emission). Controls include co-nucleofection of Cas9 RNP and ssODN (white bar) and co-nucleofection of Cas9 RNP and ssODN followed by dosing with 2,500 nM $LF_N$-Acr (− PA) at a 0-h dosing time. The values were normalized to Cas9 RNP + ssODN and are the mean ± SD of three independent replicates. The significance of $LF_N$-Acr/PA additions was determined with an unpaired, two-tailed *t*-test versus Cas9 RNP + ssODN. The differences are not statistically significant.

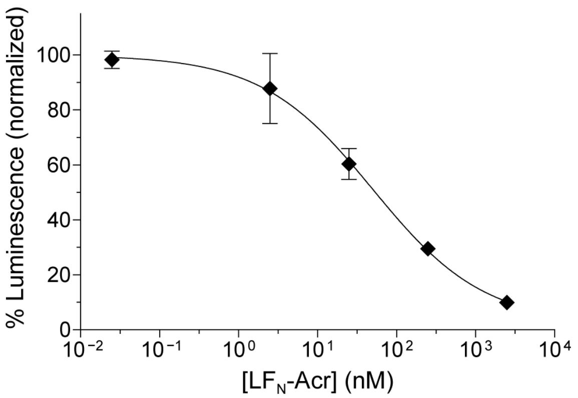


**Fig. S12.** Dose-dependent inhibition of dCas9-VPR by $LF_N$-Acr in the CRISPRa assay. dSpCas9-VPR plasmid (300 ng) was delivered via nucleofection into 7×sgRNA-CRE-NanoLuc-HEK293T cells. The cells were incubated at 37 °C and dosed with $LF_N$-Acr (0.025, 2.5, 25, 250, and 2,500 nM) and PA (20 nM) at a 0-h dosing time. After 24 h, the cells were incubated with furimazine, and their luminescence was quantified. The values were normalized to dCas9-VPR plasmid (100%, not shown) and are the mean ± SD of three independent replicates. Some error bars are too small to visualize.

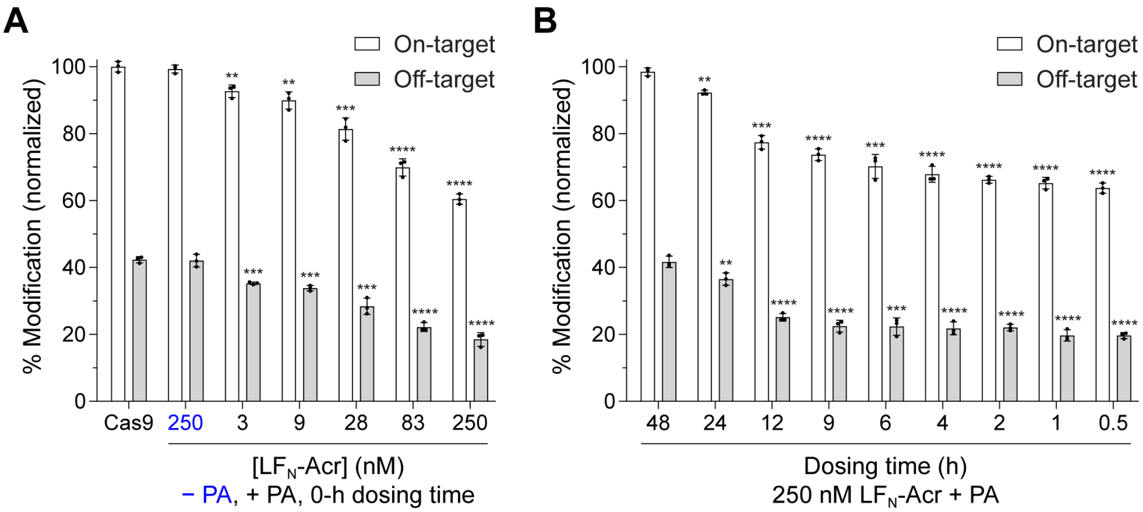


**Fig. S13.** Dose- and time-dependent inhibition of Cas9 by $LF_N$-Acr, assessed via next-generation sequencing (NGS). NLS-SpCas9-NLS RNP (80 pmol), targeting *EMX1*, was delivered via nucleofection into HEK293T cells. The cells were incubated at 37 °C and dosed with $LF_N$-Acr (3–250 nM) and PA (20 nM) at a 0-h dosing time (*A*) or dosed with $LF_N$-Acr (250 nM) and PA (20 nM), varying the dosing time from 0.5–48 h (*B*). After 72 h, genomic DNA was extracted, sequenced via NGS, and analyzed using the CRISPResso2 software pipeline to determine insertions, deletions, and substitutions (% modification) at the on-target and off-target sites. Controls include nucleofection of Cas9 RNP (panel *A*) and nucleofection of Cas9 RNP followed by dosing with 250 nM $LF_N$-Acr (− PA, panel *A*) at a 0-h dosing time. The values were normalized to Cas9 RNP (on-target) and are the mean ± SD of three independent replicates. The significance of $LF_N$-Acr/PA additions was determined with an unpaired, two-tailed *t*-test versus Cas9 RNP (on-target and off-target), where *, **, ***, and **** refer to $P \leq 0.05$, $P \leq 0.01$, $P \leq 0.001$, and $P \leq 0.0001$, respectively.

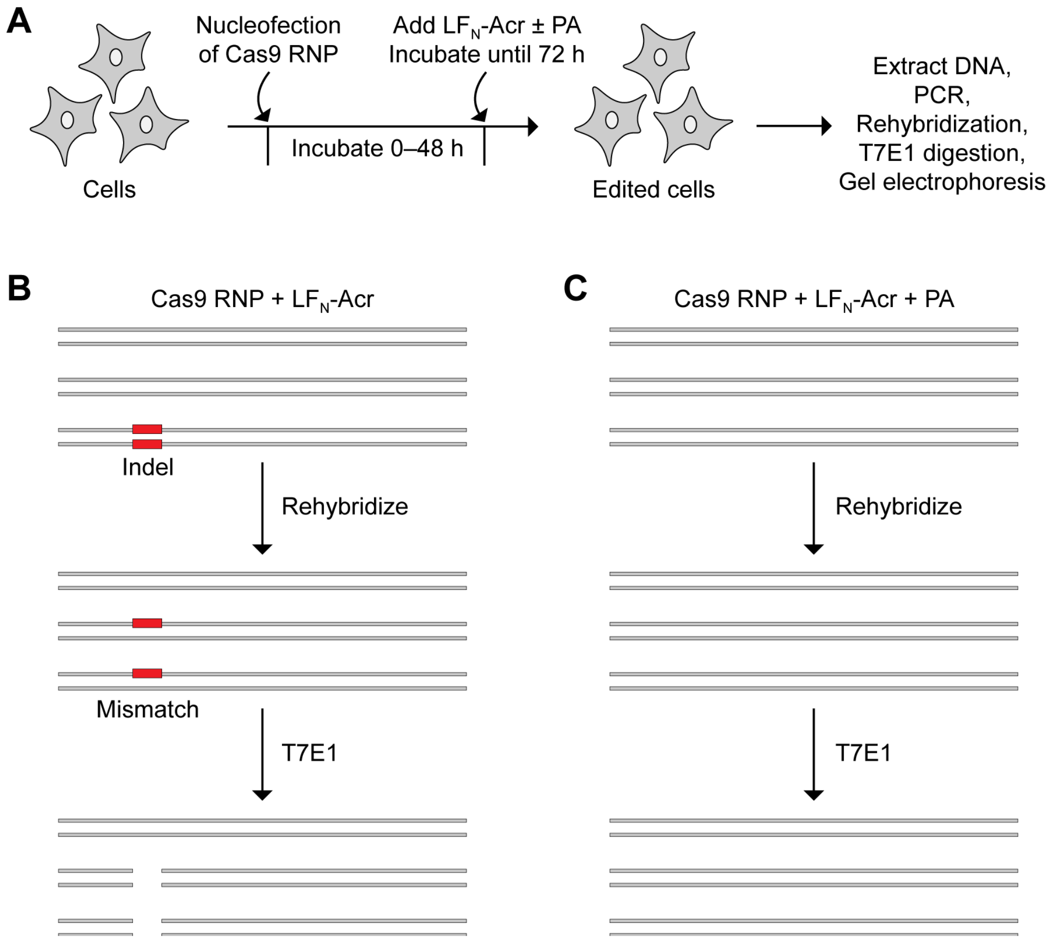


**Fig. S14.** T7 endonuclease 1 (T7E1) to assess $LF_N$-Acr delivery. (*A*) A cartoon showing the experimental set-up for the T7E1 assay. After delivering Cas9 RNP via nucleofection and incubating the cells with $LF_N$-Acr ± PA, genomic DNA is extracted from the cells and PCR amplified. After rehybridization, the amplicon is incubated with T7E1, and the cleavage products are visualized via agarose gel electrophoresis. (*B*) Because $LF_N$-Acr cannot enter cells without PA, Cas9 cleaves the DNA target, which leads to insertions and deletions (indels) that cause mismatches after rehybridization of the amplicon. T7E1 recognizes and cleaves these mismatches. (*C*) The PA-mediated delivery of $LF_N$-Acr prevents Cas9-mediated cleavage, indels, and mismatches; therefore, T7E1 does not cleave the amplicon.

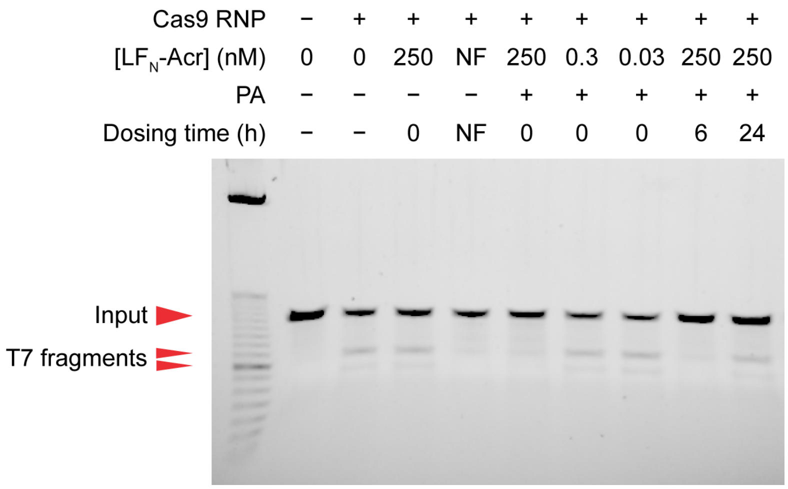


**Fig. S15.** Dose- and time-dependent inhibition of Cas9 by LF$_N$-Acr in the T7E1 assay. C-NLS-SpCas9 RNP (20 pmol), targeting *EMX1*, was delivered via nucleofection into HEK293T cells. The cells were incubated at 37 °C and dosed with LF$_N$-Acr (0.03–250 nM) and PA (20 nM) after 0–24 h (dosing time). After 72 h, genomic DNA was extracted, and the target was PCR amplified, rehybridized, incubated with T7E1, and visualized via agarose gel electrophoresis. Positive controls include nucleofection of apo-Cas9 and co-nucleofection of Cas9 RNP and 50 pmol LF$_N$-Acr (NF). Negative controls include nucleofection of Cas9 RNP and nucleofection of Cas9 RNP followed by incubation with LF$_N$-Acr (250 nM) at a 0-h dosing time. The first lane contains an E-Gel 50-bp DNA ladder with the following sizes (in base pairs): 2,500, 800, 750, 700, 650, 600, 550, 500, 450, 400, 350, 300, 250, 200, 150, 100, and 50 bp. The size (bp) of the higher-intensity bands in the ladder is underlined.

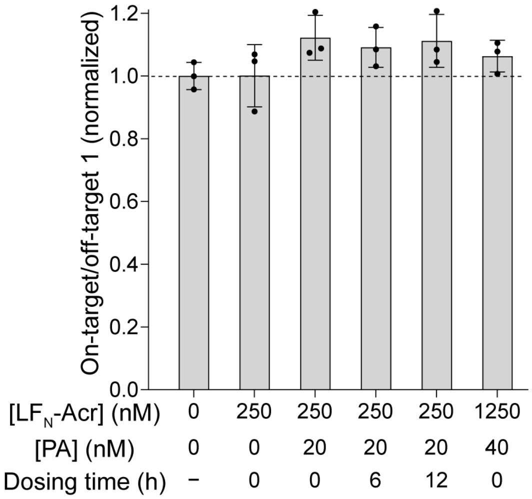


**Fig. S16.** Effect of $LF_N$-Acr delivery on base-editing specificity. ABE8e (500 ng) and sgRNA (165 ng) plasmids, targeting *HBG2*, were delivered via nucleofection into HEK293T cells. The cells were incubated at 37 °C and dosed with $LF_N$-Acr (250–1250 nM) and PA (20–40 nM) after 0–12 h (dosing time). After 72 h, genomic DNA was extracted, sequenced via NGS, and analyzed using CRISPResso2 to determine % modification at the on-target (On) and off-target (OT1, OT2, and OT3) sites. Controls include nucleofection of ABE8e-sgRNA plasmids and nucleofection of ABE8e-sgRNA plasmids followed by dosing with $LF_N$-Acr (250 nM) at a 0-h dosing time. The specificity was calculated as the on-target to off-target % modification ratio, normalizing to the on-target to off-target % modification ratio of ABE8e-sgRNA (specificity = 1). The significance of the specificity was determined with an unpaired, two-tailed *t*-test versus the specificity of ABE8e-sgRNA, where *, **, ***, and **** refer to $P \leq 0.05$, $P \leq 0.01$, $P \leq 0.001$, and $P \leq 0.0001$, respectively.

### Supplementary Tables

**Table S1.** Compilation of $IC_{50}$ and $IC_5$ values

| Inhibitor | Dosing time (h) | Assay | Cas9 | Fig. | $IC_{50}$ (nM)* | $IC_5$ (nM)* |
|---|---|---|---|---|---|---|
| $LF_N$-Acr | 0 | *GFP* | RNP | S4 | 2.9 | 0.086 |
| $LF_N$-Acr | 2 | *GFP* | RNP | 2B, S4 | 0.35 | 0.0031 |
| BRD7586 | 2 | *GFP* | RNP | 2B | — | 10,000† |
| $LF_N$-Acr | 0 | *HiBiT* | RNP | S10 | 17 | 0.044 |
| $LF_N$-Acr | 2 | *HiBiT* | RNP | S10 | 0.71 | 0.021 |
| $LF_N$-Acr | 0 | CRISPRa | Plasmid | S12 | 54 | 0.44 |

*The "Absolute $IC_{50}$, X is log(concentration)" function in GraphPad Prism software was used to fit the $LF_N$-Acr delivery data from the *GFP*-disruption, *HiBiT*-knock-in, and CRISPRa assays and determine $IC_{50}$ and $IC_5$ values.

†The $IC_5$ value for BRD7586 was determined experimentally, rather than using the "Absolute $IC_{50}$, X is log(concentration)" function, because the BRD7586 dose curve did not reach a stable baseline. BRD7586 (10 µM) reduced Cas9 activity (always normalized to 100% without Acr) from 100–95% at a 2-h dosing time (Fig. 2B). The $IC_5$ of BRD7586 is 10 µM because, at this concentration, BRD7586 inhibited Cas9 activity by 5%.

**Table S2.** Cell lines used in this study

| Cell Line | Purpose | Reference |
|---|---|---|
| NEB® 5-alpha Competent *E. coli* (High Efficiency) | Cloning the $LF_N$-Acr bacterial production plasmid | NEB |
| Rosetta 2(DE3) Competent Cells (Novagen) | Producing and purifying $LF_N$-Acr | MilliporeSigma |
| NEB® Stable Competent *E. coli* (High Efficiency) | Cloning plasmids for mammalian production | NEB |
| U2OS-EGFP.PEST | *GFP*-disruption assay | Reyon et al. (2012) (1) |
| HEK293T | *HiBiT*, T7E1, and NGS experiments | ATCC |
| 7×sgRNA-CRE-NanoLuc-HEK293T | CRISPRa assay | This work |

**Table S3.** Proteins used in this study

| Protein | Source | Reference |
|---|---|---|
| $LF_N$-Acr | Purified in-house | This work |
| Protective Antigen | Purified in-house | Pomerantsev et al. (2017) (2) |
| NLS-Cas9-NLS | Purified in-house | Lim et al. (2020) (3) |
| C-NLS-SpCas9 | GenScript | Cat. No. Z03385 |
| TEV protease | Purified in-house | Kapust et al. (2001) (4) |

**Table S4.** Plasmids used in this study

| Plasmid name | Purpose | Reference |
|---|---|---|
| pKEW336-MBP-TEV-MbCas12a-33362 | Cloning pAV14 (backbone) | Watters et al. (2018) (5) Addgene #115670 |
| pET-SUMO-$LF_N$ | Amplifying the $LF_N$ insert used to clone pAV14 | Ling et al. (2012) (6) |
| pAV14-MBP-TEV-$LF_N$-AcrIIA4 | Bacterial production and purification of $LF_N$-Acr | This work |
| pYS5-PA BH500 | Bacterial production and purification of PA | Singh et al. (1989) (7) |
| pET28a-Cas9-His | Bacterial production and purification of NLS-SpCas9-NLS | Liang et al. (2017) (8) Addgene #98158 |
| pRK793 | Bacterial production and purification of TEV protease | Kapust et al. (2001) (4) Addgene #8827 |
| Sp-dCas9-VPR | Mammalian production of dSpCas9-VPR | Chavez et al. (2015) (9) Addgene #63798 |
| ABE8e | Mammalian production of ABE8e | Richter et al. (2020) (10) Addgene #138489 |
| pHBG2-sgRNA | Mammalian production of HBG2 sgRNA | Richter et al. (2020) (10) |
| 7×CRE gRNAs | Stable sgRNA production in HEK293T cells | This work |
| CRE-NanoLuc.PEST | Stable CRE-dependent NanoLuc production in HEK293T | This work |

**Table S5.** Primer sequences for gRNA cloning and synthesis for RNP experiments

| Primer | Sequence* |
|---|---|
| GFP Fwd | TAATACGACTCACTATA**GGTGGTGCAGATGAACTTCA**GTTTTAGAGCTAGAAAT |
| GAPDH Fwd | TAATACGACTCACTATA**GGTCCAGGGGTCTTACTCCT**GTTTTAGAGCTAGAAAT |
| EMX1 Fwd | TAATACGACTCACTATA**GAGTCCGAGCAGAAGAAGAA**GTTTTAGAGCTAGAAAT |
| Universal Rev | AAAAGCACCGACTCGGTGCCACTTTTTCAAGTTGATAACGGACTAGCCTTATTTTAACTTGCT ATTTCTAGCTCTAAAAC |

*The spacer sequence is in bold typeface.

**Table S6.** Spacers of gRNA sequences encoded in plasmids

| Plasmid | Sequence |
|---|---|
| 7×CRE | GACACCCCATTGACGTCAAT,<br>ACGTCAGCTGCCAGATCCCA,<br>TGGGAGAACAGATCTGGCCT,<br>GCGCTAGCGAGCTCAGGTAC,<br>GAGAACAGATCTGGCCTCGG,<br>TGAAAGACGTCACAGTATGA,<br>AGTATGACGGCCATGGGATC |
| HBG2 | GTGGGGAAGGGGCCCCCAAG |

**Table S7.** ssODN sequence

| | |
|---|---|
| GAPDH HiBiT | TCTTCTAGGTATGACAACGAATTTGGCTACAGCAACAGGGTGGTGGACCTCATGGCCCACATGGCCT<br>CCAAGGAGGTGAGCGGCTGGCGGCTGTTCAAGAAGATTAGCTAAGACCCCTGGACCACCAGCCCCAG<br>CAAGAGCACAAGAGGAAGAGAGAGACCCTCACTGCTGGGGAGTCCCTGC |

**Table S8.** Primers used to generate amplicons for next-generation sequencing

| | Target | PAM | Forward Primer | Reverse Primer |
|---|---|---|---|---|
| EMX1 On | GAGTCCGAGCAGAAGAAGAA | GGG | ACACTCTTTCCCTACACGACGCTCTTCCGATCTNNNNCAGCTCAGCCTGAGTGTTGA | TGGAGTTCAGACGTGTGCTCTTCCGATCTCTCGTGGGTTTGTGGTTGC |
| EMX1 OT1 | GAGTTAGAGCAGAAGAAGAA | AGG | ACACTCTTTCCCTACACGACGCTCTTCCGATCTNNNNTTCTGAGGGCTGCTACCTGT | TGGAGTTCAGACGTGTGCTCTTCCGATCTGCCCAATCATTGATGCTTTT |
| HBG2 On | GTGGGGAAGGGGCCCCCAAG | AGG | ACACTCTTTCCCTACACGACGCTCTTCCGATCTNNNNGTGGAGTTTAGCCAGGGACC | TGGAGTTCAGACGTGTGCTCTTCCGATCTTGGTGGGAGAAGAAAACTAG |
| HBG2 OT1 | GGTGGGATGGGGTCCCCAAG | TGG | ACACTCTTTCCCTACACGACGCTCTTCCGATCTNNNNATGGCTGCAAATCCAAGGGT | TGGAGTTCAGACGTGTGCTCTTCCGATCTAAATGCTTCTCGGGCTCTCC |
| HBG2 OT2 | GGTAGGGAGAGGCCCCCAGA | GGG | ACACTCTTTCCCTACACGACGCTCTTCCGATCTNNNNGAGGTTGAAACTCCTCGCCA | TGGAGTTCAGACGTGTGCTCTTCCGATCTGGAATTAAGATGCAACTGAGAGTA |
| HBG2 OT3 | GGTGGGGAGCGGCCCCCCAG | TGG | ACACTCTTTCCCTACACGACGCTCTTCCGATCTNNNNGGCTGTCCCTGGTTGTCTGG | TGGAGTTCAGACGTGTGCTCTTCCGATCTCGAGCACTGAGGCCTGGTTA |

**Table S9.** Amplicon sequences

| Target | Sequence |
|---|---|
| EMX1 On | CAGCTCAGCCTGAGTGTTGAGGCCCCAGTGGCTGCTCTGGGGGCCTCCTGAGTTTCTCATCTGTGCCCCTCCCTCCCTGGCCCAGGTGAAGGTGTGGTTCCAGAACCGGAGGACAAAGTACAAACGGCAGAAGCTGGAGGAGGAAGGGCCTGAGTCCGAGCAGAAGAAGAAGGGCTCCCATCACATCAACCGGTGGCGCATTGCCACGAAGCAGGCCAATGGGGAGGACATCGATGTCACCTCCAATGACTAGGGTGGGCAACCACAAACCCACGAG |
| EMX1 OT1 | TTCTGAGGGCTGCTACCTGTACATCTGCACAAGATTGCCTTTACTCCATGCCTTTCTTCTTCTGCTCTAACTCTGACAATCTGTCTTGCCATGCCATAAGCCCCTATTCTTTCTGTAACCCCAAGATGGTATAAAAGCATCAATGATTGGGC |
| HBG2 On | GTGGAGTTTAGCCAGGGACCGTTTCAGACAGATATTTGCATTGAGATAGTGTGGGGAAGGGGCCCCCAAGAGGATACTGCTGCTTAATTTTTTTTATAGCCTTTGCCTTGTTCCGATTCAGTCATTCCAATTTTTCTCTAATTTATTCTTCCCTTTAGCTAGTTTTCTTCTCCCACCA |
| HBG2 OT1 | ATGGCTGCAAATCCAAGGGTGCACACTGGGGTGTCCTTGGCCTGAGTGCCAGCCTGCAGAGAGCAGGAGTGGGCCACTTGGGGACCCCATCCCACCAGGAAGCCTGAGAAAGCCCAGGGTGAGCCCGGCCCTGTGGAATTCTGGAGAGCCCGAGAAGCATTT |
| HBG2 OT2 | GAGGTTGAAACTCCTCGCCACAGTCCACTCTCCATCTCCATCCCTACCCCATCAATAAGCCCTTCCCCCACCATCTCAAGACACTGGAATGCAGGTAGGGAGAGGCCCCCAGAGGGAGTATTTGAGAATTTGCTCCCCTGTCCCTTGCCTCAACTACTCTCAGTTGCATCTTAATTCC |
| HBG2 OT3 | GGCTGTCCCTGGTTGTCTGGTACCTGGCTTTGGAGCAATGAGGGCTGGTGGGGAGCGGCCCCCCAGTGGGGCCGTCTGATTAGCGCAGGGCTGCCGTGTGGACCAATCAGCTGCTCAGGAAGGGGCCGGCCCCTAACCAGGCCTCAGTGCTCG |

#### Plasmid Sequences

**Sequence S1:** pAV14-MBP-TEV-LF$_N$-AcrIIA4

The sequence encoding 10×His-MBP-TEV-LF$_N$-G$_4$CG$_4$S-AcrIIA4-GS-NLS is in uppercase, and its domains are underlined differently: 10×His-MBP tag, TEV site, LF$_N$, G$_4$CG$_4$S linker, **AcrIIA4**, GS linker, SV40 NLS.

tcccccggccacggggcctgccaccatacccacgccgaaacaagcgctcatgagcccgaagtggcgagccc
gatcttccccatcggtgatgtcggcgatataggcgccagcaaccgcacctgtggcgccggtgatgccggcc
acgatgcgtccggcgtagaggatcgagatctcgatcccgcgaaattaatacgactcactatagggagacca
caacggtttccctctagtgccggctccggagagctctttaattaagcggccgccctgcaggactcgagttc
tagaaataattttgtttaactttaagaaggagatatacatATGAAATCTTCTCACCATCACCATCACCATC
ACCATCACCATGGTTCTTCTATGAAAATCGAAGAAGGTAAACTGGTAATCTGGATTAACGGCGATAAAGGC
TATAACGGTCTCGCTGAAGTCGGTAAGAAATTCGAGAAAGATACCGGAATTAAAGTCACCGTTGAGCATCC
GGATAAACTGGAAGAGAAATTCCCACAGGTTGCGGCAACTGGCGATGGCCCTGACATTATCTTCTGGGCAC
ACGACCGCTTTGGTGGCTACGCTCAATCTGGCCTGTTGGCTGAAATCACCCCGGACAAAGCGTTCCAGGAC
AAGCTGTATCCGTTTACCTGGGATGCCGTACGTTACAACGGCAAGCTGATTGCTTACCCGATCGCTGTTGA
AGCGTTATCGCTGATTTATAACAAAGATCTGCTGCCGAACCCGCCAAAAACCTGGGAAGAGATCCCGGCGC
TGGATAAAGAACTGAAAGCGAAAGGTAAGAGCGCGCTGATGTTCAACCTGCAAGAACCGTACTTCACCTGG
CCGCTGATTGCTGCTGACGGGGGTTATGCGTTCAAGTATGAAAACGGCAAGTACGACATTAAAGACGTGGG
CGTGGATAACGCTGGCGCGAAAGCGGGTCTGACCTTCCTGGTTGACCTGATTAAAAACAAACACATGAATG
CAGACACCGATTACTCCATCGCAGAAGCTGCCTTTAATAAAGGCGAAACAGCGATGACCATCAACGGCCCG
TGGGCATGGTCCAACATCGACACCAGCAAAGTGAATTATGGTGTAACGGTACTGCCGACCTTCAAGGGTCA
ACCATCCAAACCGTTCGTTGGCGTGCTGAGCGCAGGTATTAACGCCGCCAGTCCGAACAAAGAGCTGGCAA
AAGAGTTCCTCGAAAACTATCTGCTGACTGATGAAGGTCTGGAAGCGGTTAATAAAGACAAACCGCTGGGT
GCCGTAGCGCTGAAGTCTTACGAGGAAGAGTTGGCGAAAGATCCACGTATTGCCGCCACTATGGAAAACGC
CCAGAAAGGTGAAATCATGCCGAACATCCCGCAGATGTCCGCTTTCTGGTATGCCGTGCGTACTGCGGTGA
TCAACGCCGCCAGCGGTCGTCAGACTGTCGATGAAGCCCTGAAAGACGCGCAGACTAATTCGAGCTCGAAC
AACAACAACAATAACAATAACAACAACCTCGGGATCGAGGAAAACCTGTACTTCCAATCCAATGCAGCGGG
CGGTCATGGTGATGTAGGTATGCACGTAAAAGAGAAAGAGAAAAATAAAGATGAGAATAAGAGAAAAGATG
AAGAACGAAATAAAACACAGGAAGAGCATTTAAAGGAAATCATGAAACACATTGTAAAAATAGAAGTAAAA
GGGGAGGAAGCTGTTAAAAAAGAGGCAGCAGAAAAGCTACTTGAGAAAGTACCATCTGATGTTTTAGAGAT
GTATAAAGCAATTGGAGGAAAGATATATATTGTGGATGGTGATATTACAAAACATATATCTTTAGAAGCAT
TATCTGAAGATAAGAAAAAAATAAAAGACATTTATGGGAAAGATGCTTTATTACATGAACATTATGTATAT
GCAAAAGAAGGATATGAACCCGTACTTGTAATCCAATCTTCGGAAGATTATGTAGAAAATACTGAAAAGGC
ACTGAACGTTTATTATGAAATAGGTAAGATATTATCAAGGGATATTTTAAGTAAAATTAATCAACCATATC
AGAAATTTTTAGATGTATTAAATACCATTAAAAATGCATCTGATTCAGATGGACAAGATCTTTTATTTACT
AATCAGCTTAAGGAACATCCCACAGACTTTTCTGTAGAATTCTTGGAACAAAATAGCAATGAGGTACAAGA
AGTATTTGCGAAAGCTTTTGCATATTATATCGAGCCACAGCATCGTGATGTTTTACAGCTTTATGCACCGG
AAGCTTTTAATTACATGGATAAATTTAACGAACAAGAAATAAATCTATCCTTGGAAGAACTTAAAGATCAA
CGGGGCGGTGGTGGTTGTGGTGGGGGCGGCAGC**ATGAATATTAATGACTTAATTAGAGAAATCAAAAACAA**
**AGATTACACAGTGAAATTGAGTGGTACGGATAGCAATAGTATCACACAGCTAATTATTCGCGTTAATAATG**
**ATGGCAACGAGTATGTAATTTCTGAAAGTGAAAATGAATCAATCGTTGAAAAATTCATCTCTGCATTCAAA**
**AACGGTTGGAATCAAGAATACGAGGATGAAGAAGAATTTTATAATGACATGCAAACAATCACCTTAAAAAG**
**TGAGTTGAAC**GGGTCACCTAAGAAAAAACGAAAAGTTtaataaatattggaagtggataacggatccgcga
tcgcggcgcgccacctggtggccggccggtaccacgcgtgcgcgctgatccggctgctaacaaagcccgaa

aggaagctgagttggctgctgccaccgctgagcaataactagcataaccccttggggcctctaaacgggtc
ttgaggggttttttgctgaaaggaggaactatatccggatatccacaggacgggtgtggtcgccatgatcg
cgtagtcgatagtggctccaagtagcgaagcgagcaggactgggcggcggccaaagcggtcggacagtgct
ccgagaacgggtgcgcatagaaattgcatcaacgcatatagcgctagcagcacgccatagtgactggcgat
gctgtcggaatggacgatatcccgcaagaggcccggcagtaccggcataaccaagcctatgcctacagcat
ccagggtgacggtgccgaggatgacgatgagcgcattgttagatttcatacacggtgcctgactgcgttag
caatttaactgtgataaactaccgcattaaagcttatcgatgataagctgtcaaacatgagaattcttgaa
gacgaaagggcctcgtgatacgcctatttttataggttaatgtcatgataataatggtttcttagacgtca
ggtggcacttttcggggaaatgtgcgcggaacccctatttgtttatttttctaaatacattcaaatatgta
tccgctcatgagacaataaccctgataaatgcttcaataacattgaaaaaggaagagtatgagtattcaac
atttccgtgtcgcccttattcccttttttgcggcattttgccttcctgtttttgctcacccagaaacgctg
gtgaaagtaaaagatgctgaagatcagttgggtgcacgagtgggttacatcgaactggatctcaacagcgg
taagatccttgagagttttcgccccgaagaacgttttccaatgatgagcacttttaaagttctgctatgtg
gcgcggtattatcccgtgttgacgccgggcaagagcaactcggtcgccgcatacactattctcagaatgac
ttggttgagtactcaccagtcacagaaaagcatcttacggatggcatgacagtaagagaattatgcagtgc
tgccataaccatgagtgataacactgcggccaacttacttctgacaacgatcggaggaccgaaggagctaa
ccgcttttttgcacaacatgggggatcatgtaactcgccttgatcgttgggaaccggagctgaatgaagcc
ataccaaacgacgagcgtgacaccacgatgcctgcagcaatggcaacaacgttgcgcaaactattaactgg
cgaactacttactctagcttcccggcaacaattaatagactggatggaggcggataaagttgcaggaccac
ttctgcgctcggcccttccggctggctggtttattgctgataaatctggagccggtgagcgtgggtctcgc
ggtatcattgcagcactggggccagatggtaagccctcccgtatcgtagttatctacacgacggggagtca
ggcaactatggatgaacgaaatagacagatcgctgagataggtgcctcactgattaagcattggtaactgt
cagaccaagtttactcatatatactttagattgatttaaaacttcatttttaatttaaaaggatctaggtg
aagatcctttttgataatctcatgaccaaaatcccttaacgtgagttttcgttccactgagcgtcagaccc
cgtagaaaagatcaaaggatcttcttgagatcctttttttctgcgcgtaatctgctgcttgcaaacaaaaa
aaccaccgctaccagcggtggtttgtttgccggatcaagagctaccaactctttttccgaaggtaactggc
ttcagcagagcgcagataccaaatactgtccttctagtgtagccgtagttaggccaccacttcaagaactc
tgtagcaccgcctacatacctcgctctgctaatcctgttaccagtggctgctgccagtggcgataagtcgt
gtcttaccgggttggactcaagacgatagttaccggataaggcgcagcggtcgggctgaacggggggttcg
tgcacacagcccagcttggagcgaacgacctacaccgaactgagatacctacagcgtgagctatgagaaag
cgccacgcttcccgaagggagaaaggcggacaggtatccggtaagcggcagggtcggaacaggagagcgca
cgagggagcttccagggggaaacgcctggtatctttatagtcctgtcgggtttcgccacctctgacttgag
cgtcgatttttgtgatgctcgtcaggggggcggagcctatggaaaaacgccagcaacgcggcctttttacg
gttcctggccttttgctggccttttgctcacatgttctttcctgcgttatcccctgattctgtggataacc
gtattaccgcctttgagtgagctgataccgctcgccgcagccgaacgaccgagcgcagcgagtcagtgagc
gaggaagcggaagagcgcctgatgcggtattttctccttacgcatctgtgcggtatttcacaccgcaatgg
tgcactctcagtacaatctgctctgatgccgcatagttaagccagtatacactccgctatcgctacgtgac
tgggtcatggctgcgccccgacacccgccaacacccgctgacgcgccctgacgggcttgtctgctcccggc
atccgcttacagacaagctgtgaccgtctccgggagctgcatgtgtcagaggttttcaccgtcatcaccga
aacgcgcgaggcagctgcggtaaagctcatcagcgtggtcgtgaagcgattcacagatgtctgcctgttca
tccgcgtccagctcgttgagtttctccagaagcgttaatgtctggcttctgataaagcgggccatgttaag
ggcggttttttcctgtttggtcactgatgcctccgtgtaagggggatttctgttcatgggggtaatgatac
cgatgaaacgagagaggatgctcacgatacgggttactgatgatgaacatgcccggttactggaacgttgt
gagggtaaacaactggcggtatggatgcggcgggaccagagaaaaatcactcagggtcaatgccagcgctt
cgttaatacagatgtaggtgttccacagggtagccagcagcatcctgcgatgcagatccggaacataatgg

tgcagggcgctgacttccgcgtttccagactttacgaaacacggaaaccgaagaccattcatgttgttgct
caggtcgcagacgttttgcagcagcagtcgcttcacgttcgctcgcgtatcggtgattcattctgctaacc
agtaaggcaaccccgccagcctagccgggtcctcaacgacaggagcacgatcatgcgcacccgtggccagg
acccaacgctgcccgagatgcgccgcgtgcggctgctggagatggcggacgcgatggatatgttctgccaa
gggttggtttgcgcattcacagttctccgcaagaattgattggctccaattcttggagtggtgaatccgtt
agcgaggtgccgccggcttccattcaggtcgaggtggcccggctccatgcaccgcgacgcaacgcggggag
gcagacaaggtatagggcggcgcctacaatccatgccaacccgttccatgtgctcgccgaggcggcataaa
tcgccgtgacgatcagcggtccaatgatcgaagttaggctggtaagagccgcgagcgatccttgaagctgt
ccctgatggtcgtcatctacctgcctggacagcatggcctgcaacgcgggcatcccgatgccgccggaagc
gagaagaatcataatggggaaggccatccagcctcgcgtcgcgaacgccagcaagacgtagcccagcgcgt
cggccgccatgccggcgataatggcctgcttctcgccgaaacgtttggtggcgggaccagtgacgaaggct
tgagcgagggcgtgcaagattccgaataccgcaagcgacaggccgatcatcgtcgcgctccagcgaaagcg
gtcctcgccgaaaatgacccagagcgctgccggcacctgtcctacgagttgcatgataaagaagacagtca
taagtgcggcgacgatagtcatgccccgcgcccaccggaaggagctgactgggttgaaggctctcaagggc
atcggtcgacgctctcccttatgcgactcctgcattaggaagcagcccagtagtaggttgaggccgttgag
caccgccgccgcaaggaatggtgcatgcaaggagatggcgcccaacag

**Sequence S2:** 7×CRE gRNAs (this work, stably expressed in 7×sgRNA-CRE-NanoLuc-HEK293T cells)

ttgatgcctggcagttccctactctcgcgttaacgctagcatggatgttttcccagtcacgacgttgtaaa
acgacggccagtcttaagcgtctcatggcctgaccccggaccaagtggtggctatgagggcctatttccca
tgattccttcatatttgcatatacgatacaaggctgttagagagataattggaattaatttgactgtaaac
acaaagatattagtacaaaatacgtgacgtagaaagtaataatttcttgggtagtttgcagttttaaaatt
atgttttaaaatggactatcatatgcttaccgtaacttgaaagtatttcgatttcttggctttatatatct
tgtggaaaggacgaaacaccgacaccccattgacgtcaatgttttagagctagaaatagcaagttaaaata
aggctagtccgttatcaacttgaaaaagtggcaccgagtcggtgcttttttgttttagagctagaaatagc
aagttacatggagggcctatttcccatgattccttcatatttgcatatacgatacaaggctgttagagaga
taattggaattaatttgactgtaaacacaaagatattagtacaaaatacgtgacgtagaaagtaataattt
cttgggtagtttgcagttttaaagttatgttttaaaatggactatcatatgcttaccgtaacttgaaagta
tttcgatttcttggctttatatatcttgtggaaaggacgaaacaccacgtcagctgccagatcccagtttt
agagctagaaatagcaagttaaaataaggctagtccgttatcaacttgaaaaagtggcaccgagtcggtgc
ttttttgttttagagctagaaatagcaagttaggacgagggcctatttcccatgattccttcatatttgca
tatacgatacaaggctgttagagagataattggaattaatttgactgtaaacacaaagatattagtacaaa
atacgtgacgtagaaagtaataatttcttgggtagtttgcagttttaaaattatgttttaaaatggactat
catatgcttaccgtaacttgaaagtatttcgatttcttggctttatatatcttgtggaaaggacgaaacac
ctgggagaacagatctggcctgttttagagctagaaatagcaagttaaaataaggctagtccgttatcaac
ttgaaaaagtggcaccgagtcggtgcttttttgttttagagctagaaatagcaagttaccaggagggccta
tttcccatgattccttcatatttgcatatacgatacaaggctgttagagagataattggaattaatttgac
tgtaaacacaaagatattagtacaaaatacgtgacgtagaaagtaataatttcttgggtagtttgcagttt
taaaattatgttttaaaatggactatcatatgcttaccgtaacttgaaagtatttcgatttcttggcttta
tatatcttgtggaaaggacgaaacaccgcgctagcgagctcaggtacgttttagagctagaaatagcaagt
taaaataaggctagtccgttatcaacttgaaaaagtggcaccgagtcggtgcttttttgttttagagctag
aaatagcaagttatgttgagggcctatttcccatgattccttcatatttgcatatacgatacaaggctgtt
agagagataattggaattaatttgactgtaaacacaaagatattagtacaaaatacgtgacgtagaaagta
ataatttcttgggtagtttgcagttttaaaattatgttttaaaatggactatcatatgcttaccgtaactt
gaaagtatttcgatttcttggctttatatatcttgtggaaaggacgaaacaccgagaacagatctggcctc
gggttttagagctagaaatagcaagttaaaataaggctagtccgttatcaacttgaaaaagtggcaccgag
tcggtgcttttttgttttagagctagaaatagcaagttatgcagagggcctatttcccatgattccttcat
atttgcatatacgatacaaggctgttagagagataattggaattaatttgactgtaaacacaaagatatta
gtacaaaatacgtgacgtagaaagtaataatttcttgggtagtttgcagttttaaaattatgttttaaaat
ggactatcatatgcttaccgtaacttgaaagtatttcgatttcttggctttatatatcttgtggaaaggac
gaaacacctgaaagacgtcacagtatgagttttagagctagaaatagcaagttaaaataaggctagtccgt
tatcaacttgaaaaagtggcaccgagtcggtgcttttttgttttagagctagaaatagcaagttacggtga
gggcctatttcccatgattccttcatatttgcatatacgatacaaggctgttagagagataattggaatta
atttgactgtaaacacaaagatattagtacaaaatacgtgacgtagaaagtaataatttcttgggtagttt
gcagttttaaaattatgttttaaaatggactatcatatgcttaccgtaacttgaaagtatttcgatttctt
ggctttatatatcttgtggaaaggacgaaacaccagtatgacggccatgggatcgttttagagctagaaat
agcaagttaaaataaggctagtccgttatcaacttgaaaaagtggcaccgagtcggtgcttttttgtttta
gagctagaaatagcaagttagaaacggtgcagcggctgttgccggtgctgtgccaggaccatggcctgacc
ccggaccaagtggtggctatcgagacgtctagaccagccaggacagaaatgcctcgacttcgctgctaccc
aaggttgccgggtgacgcacaccgtggaaacggatgaaggcacgaacccagtggacataagcctgttcggt
tcgtaagctgtaatgcaagtagcgtatgcgctcacgcaactggtccagaaccttgaccgaacgcagcggtg

gtaacggcgcagtggcggttttcatggcttgttatgactgtttttttggggtacagtctatgcctcgggca
tccaagcagcaagcgcgttacgccgtgggtcgatgtttgatgttatggagcagcaacgatgttacgcagca
gggcagtcgccctaaaacaaagttaaacattatgagggaagcggtgatcgccgaagtatcgactcaactat
cagaggtagttggcgtcatcgagcgccatctcgaaccgacgttgctggccgtacatttgtacggctccgca
gtggatggcggcctgaagccacacagtgatattgatttgctggttacggtgaccgtaaggcttgatgaaac
aacgcggcgagctttgatcaacgaccttttggaaacttcggcttcccctggagagagcgagattctccgcg
ctgtagaagtcaccattgttgtgcacgacgacatcattccgtggcgttatccagctaagcgcgaactgcaa
tttggagaatggcagcgcaatgacattcttgcaggtatcttcgagccagccacgatcgacattgatctggc
tatcttgctgacaaaagcaagagaacatagcgttgccttggtaggtccagcggcggaggaactctttgatc
cggttcctgaacaggatctatttgaggcgctaaatgaaaccttaacgctatggaactcgccgcccgactgg
gctggcgatgagcgaaatgtagtgcttacgttgtcccgcatttggtacagcgcagtaaccggcaaaatcgc
gccgaaggatgtcgctgccgactgggcaatggagcgcctgccggcccagtatcagcccgtcatacttgaag
ctagacaggcttatcttggacaagaagaagatcgcttggcctcgcgcgcagatcagttggaagaatttgtc
cactacgtgaaaggcgagatcaccaaggtagtcggcaaataaccctcgagccacccatgaccaaaatccct
taacgtgagttacgcgtcgttccactgagcgtcagaccccgtagaaaagatcaaaggatcttcttgagatc
ctttttttctgcgcgtaatctgctgcttgcaaacaaaaaaaccaccgctaccagcggtggtttgtttgccg
gatcaagagctaccaactctttttccgaaggtaactggcttcagcagagcgcagataccaaatactgtcct
tctagtgtagccgtagttaggccaccacttcaagaactctgtagcaccgcctacatacctcgctctgctaa
tcctgttaccagtggctgctgccagtggcgataagtcgtgtcttaccgggttggactcaagacgatagtta
ccggataaggcgcagcggtcgggctgaacggggggttcgtgcacacagcccagcttggagcgaacgaccta
caccgaactgagatacctacagcgtgagcattgagaaagcgccacgcttcccgaagggagaaaggcggaca
ggtatccggtaagcggcagggtcggaacaggagagcgcacgagggagcttccagggggaaacgcctggtat
ctttatagtcctgtcgggtttcgccacctctgacttgagcgtcgatttttgtgatgctcgtcaggggggcg
gagcctatggaaaaacgccagcaacgcggcctttttacggttcctggccttttgctggccttttgctcaca
tgttctttcctgcgttatcccctgattctgtggataaccgtattaccgcctttgagtgagctgataccgct
cgccgcagccgaacgaccgagcgcagcgagtcagtgagcgaggaagcggaagagcgcccaatacgcaaacc
gcctctccccgcgcgttggccgattcattaatgcagctggcacgacaggtttcccgactggaaagcgggca
gtgagcgcaacgcaattaatacgcgtaccgctagccaggaagagtttgtagaaacgcaaaaaggccatccg
tcaggatggccttctgcttagtttgatgcctggcagtttatggcgggcgtcctgcccgccaccctccgggc
cgttgcttcacaacgttcaaatccgctcccggcggatttgtcctactcaggagagcgttcaccgacaaaca
acagataaaacgaaaggcccagtcttccgactgagcctttcgttttat

**Sequence S3:** CRE-NanoLuc-PEST (this work, stably expressed in 7×sgRNA-CRE-NanoLuc-HEK293T cells)

```
cgaacgaccgagcgcagcgagtcagtgagcgaggaagcggaagagcgcccaatacgcaaaccgcctctccc
cgcgcgttggccgattcattaatgcagctggcacgacaggtttcccgactggaaagcgggcagtgagcgca
acgcaattaatgtgagttagctcactcattaggcaccccaggctttacactttatgcttccggctcgtatg
ttgtgtggaattgtgagcggataacaatttcacacaggaaacagctatgaccatgattacgccaagcgcgc
aattaaccctcactaaagggaacaaaagctggagctgcaagcttaatgtagtcttatgcaatactcttgta
gtcttgcaacatggtaacgatgagttagcaacatgccttacaaggagagaaaaagcaccgtgcatgccgat
tggtggaagtaaggtggtacgatcgtgccttattaggaaggcaacagacgggtctgacatggattggacga
accactgaattgccgcattgcagagatattgtatttaagtgcctagctcgatacataaacgggtctctctg
gttagaccagatctgagcctgggagctctctggctaactagggaacccactgcttaagcctcaataaagct
tgccttgagtgcttcaagtagtgtgtgcccgtctgttgtgtgactctggtaactagagatccctcagaccc
ttttagtcagtgtggaaaatctctagcagtggcgcccgaacagggacttgaaagcgaaagggaaaccagag
gagctctctcgacgcaggactcggcttgctgaagcgcgcacggcaagaggcgaggggcggcgactggtgag
tacgccaaaaattttgactagcggaggctagaaggagagagatgggtgcgagagcgtcagtattaagcggg
ggagaattagatcgcgatgggaaaaaattcggttaaggccagggggaaagaaaaaatataaattaaaacat
atagtatgggcaagcagggagctagaacgattcgcagttaatcctggcctgttagaaacatcagaaggctg
tagacaaatactgggacagctacaaccatcccttcagacaggatcagaagaacttagatcattatataata
cagtagcaaccctctattgtgtgcatcaaaggatagagataaaagacaccaaggaagctttagacaagata
gaggaagagcaaaacaaaagtaagaccaccgcacagcaagcggccgctgatcttcagacctggaggaggag
atatgagggacaattggagaagtgaattatataaatataaagtagtaaaaattgaaccattaggagtagca
cccaccaaggcaaagagaagagtggtgcagagagaaaaaagagcagtgggaataggagctttgttccttgg
gttcttgggagcagcaggaagcactatgggcgcagcgtcaatgacgctgacggtacaggccagacaattat
tgtctggtatagtgcagcagcagaacaatttgctgagggctattgaggcgcaacagcatctgttgcaactc
acagtctggggcatcaagcagctccaggcaagaatcctggctgtggaaagatacctaaaggatcaacagct
cctggggatttggggttgctctggaaaactcatttgcaccactgctgtgccttggaatgctagttggagta
ataaatctctggaacagatttggaatcacacgacctggatggagtgggacagagaaattaacaattacaca
agcttaatacactccttaattgaagaatcgcaaaaccagcaagaaaagaatgaacaagaattattggaatt
agataaatgggcaagtttgtggaattggtttaacataacaaattggctgtggtatataaaattattcataa
tgatagtaggaggcttggtaggtttaagaatagtttttgctgtactttctatagtgaatagagttaggcag
ggatattcaccattatcgtttcagacccacctcccaaccccgaggggacccctcaggcgtctcatggcct
gtggctatcgagacgggcctaactggccggtacctgagctcgctagcgcaccagacagtgacgtcagctgc
cagatcccatggccgtcatactgtgacgtctttcagacaccccattgacgtcaatgggagaacagatctgg
cctcggcggccaagcttagacactagagggtatataatggaagctcgacttccagcttggcaatccggtac
tgttggtaaacgccggcggccaccatggtcttcacactcgaagatttcgttggggactggcgacagacagc
cggctacaacctggaccaagtccttgaacagggaggtgtgtccagtttgtttcagaatctcggggtgtccg
taactccgatccaaaggattgtcctgagcggtgaaaatgggctgaagatcgacatccatgtcatcatcccg
tatgaaggtctgagcggcgaccaaatgggccagatcgaaaaaatttttaaggtggtgtaccctgtggatga
tcatcactttaaggtgatcctgcactatggcacactggtaatcgacggggttacgccgaacatgatcgact
atttcggacggccgtatgaaggcatcgccgtgttcgacggcaaaaagatcactgtaacagggaccctgtgg
aacggcaacaaaattatcgacgagcgcctgatcaaccccgacggctccctgctgttccgagtaaccatcaa
cggagtgaccggctggcggctgtgcgaacgcattctggcgaattctcacggcttccctcccgaggtggagg
agcaggccgccggcaccctgcccatgagctgcgcccaggagagcggcatggatagacaccctgctgcttgc
gccagcgccaggatcaacgtctaatgtacaagtaaactagtaagcttggcgtaactagatcttgagacaaa
tggcagtattcatccacaattttaaaagaaaaggggggattggggggtacagtgcaggggaaagaatagta
```

gacataatagcaacagacatacaaactaaagaattacaaaaacaaattacaaaaattcaaaattttcgggt
ttattacagggacagcagagatccactttgggctcgagggggcccgggtgcaaagatggataaagttttaa
acagagaggaatctttgcagctaatggaccttctaggtcttgaaaggagtgggaattggctccggtgcccg
tcagtgggcagagcgcacatcgcccacagtccccgagaagttggggggaggggtcggcaattgatccggtg
cctagagaaggtggcgcggggtaaactgggaaagtgatgtcgtgtactggctccgcctttttcccgagggt
gggggagaaccgtatataagtgcagtagtcgccgtgaacgttctttttcgcaacgggtttgccgccagaac
acaggtaagtgccgtgtgtggttcccgcgggcctggcctctttacgggttatggcccttgcgtgccttgaa
ttacttccacctggctgcagtacgtgattcttgatcccgagcttcgggttggaagtgggtgggagagttcg
aggccttgcgcttaaggagccccttcgcctcgtgcttgagttgaggcctggcctgggcgctggggccgccg
cgtgcgaatctggtggcaccttcgcgcctgtctcgctgctttcgataagtctctagccatttaaaattttt
gatgacctgctgcgacgctttttttctggcaagatagtcttgtaaatgcgggccaagatctgcacactggt
atttcggtttttggggccgcgggcggcgacggggcccgtgcgtcccagcgcacatgttcggcgaggcgggg
cctgcgagcgcggccaccgagaatcggacgggggtagtctcaagctggccggcctgctctggtgcctggcc
tcgcgccgccgtgtatcgccccgccctgggcggcaaggctggcccggtcggcaccagttgcgtgagcggaa
agatggccgcttcccggccctgctgcagggagctcaaaatggaggacgcggcgctcgggagagcgggcggg
tgagtcacccacacaaaggaaaagggcctttccgtcctcagccgtcgcttcatgtgactccacggagtacc
gggcgccgtccaggcacctcgattagttctcgagcttttggagtacgtcgtctttaggttggggggagggg
ttttatgcgatggagtttccccacactgagtgggtggagactgaagttaggccagcttggcacttgatgta
attctccttggaatttgccctttttgagtttggatcttggttcattctcaagcctcagacagtggttcaaa
gtttttttcttccatttcaggtgtcgtgacgtacggccaccatgaccgagtacaagcccacggtgcgcctc
gccacccgcgacgacgtccccagggccgtacgcaccctcgccgccgcgttcgccgactaccccgccacgcg
ccacaccgtcgatccggaccgccacatcgagcgggtcaccgagctgcaagaactcttcctcacgcgcgtcg
ggctcgacatcggcaaggtgtgggtcgcggacgacggcgccgccgtggcggtctggaccacgccggagagc
gtcgaagcgggggcggtgttcgccgagatcggcccgcgcatggccgagttgagcggttcccggctggccgc
gcagcaacagatggaaggcctcctggcgccgcaccggcccaaggagcccgcgtggttcctggccaccgtcg
gagtctcgcccgaccaccagggcaagggtctgggcagcgccgtcgtgctccccggagtggaggcggccgag
cgcgccggggtgcccgccttcctggagacctccgcgccccgcaacctccccttctacgagcggctcggctt
caccgtcaccgccgacgtcgaggtgcccgaaggaccgcgcacctggtgcatgacccgcaagcccggtgcct
gaacgcgttaagtcgacaatcaacctctggattacaaaatttgtgaaagattgactggtattcttaactat
gttgctccttttacgctatgtggatacgctgctttaatgcctttgtatcatgctattgcttcccgtatggc
tttcattttctcctccttgtataaatcctggttgctgtctctttatgaggagttgtggcccgttgtcaggc
aacgtggcgtggtgtgcactgtgtttgctgacgcaacccccactggttggggcattgccaccacctgtcag
ctcctttccgggactttcgctttccccctccctattgccacggcggaactcatcgccgcctgccttgcccg
ctgctggacaggggctcggctgttgggcactgacaattccgtggtgttgtcggggaaatcatcgtcctttc
cttggctgctcgcctgtgttgccacctggattctgcgcgggacgtccttctgctacgtcccttcggccctc
aatccagcggaccttccttcccgcggcctgctgccggctctgcggcctcttccgcgtcttcgccttcgccc
tcagacgagtcggatctccctttgggccgcctccccgcgtcgactttaagaccaatgacttacaaggcagc
tgtagatcttagccactttttaaaagaaaaggggggactggaagggctaattcactcccaacgaagacaag
atctgctttttgcttgtactgggtctctctggttagaccagatctgagcctgggagctctctggctaacta
gggaacccactgcttaagcctcaataaagcttgccttgagtgcttcaagtagtgtgtgcccgtctgttgtg
tgactctggtaactagagatccctcagacccttttagtcagtgtggaaaatctctagcagtacgtatagta
gttcatgtcatcttattattcagtatttataacttgcaaagaaatgaatatcagagagtgagaggaacttg
tttattgcagcttataatggttacaaataaagcaatagcatcacaaatttcacaaataaagcatttttttc
actgcattctagttgtggtttgtccaaactcatcaatgtatcttatcatgtctggctctagctatcccgcc
cctaactccgcccatcccgcccctaactccgcccagttccgcccattctccgccccatggctgactaattt

tttttatttatgcagaggccgaggccgcctcggcctctgagctattccagaagtagtgaggaggcttttt
ggaggcctagggacgtacccaattcgccctatagtgagtcgtattacgcgcgctcactggccgtcgtttta
caacgtcgtgactgggaaaaccctggcgttacccaacttaatcgccttgcagcacatccccctttcgccag
ctggcgtaatagcgaagaggcccgcaccgatcgcccttcccaacagttgcgcagcctgaatggcgaatggg
acgcgccctgtagcggcgcattaagcgcggcgggtgtggtggttacgcgcagcgtgaccgctacacttgcc
agcgccctagcgcccgctcctttcgctttcttcccttcctttctcgccacgttcgccggctttccccgtca
agctctaaatcgggggctccctttagggttccgatttagtgctttacggcacctcgaccccaaaaaacttg
attagggtgatggttcacgtagtgggccatcgccctgatagacggtttttcgccctttgacgttggagtcc
acgttctttaatagtggactcttgttccaaactggaacaacactcaaccctatctcggtctattcttttga
tttataagggattttgccgatttcggcctattggttaaaaaatgagctgatttaacaaaaatttaacgcga
attttaacaaaatattaacgcttacaatttaggtggcacttttcggggaaatgtgcgcggaacccctattt
gtttatttttctaaatacattcaaatatgtatccgctcatgagacaataaccctgataaatgcttcaataa
tattgaaaaaggaagagtatgagtattcaacatttccgtgtcgcccttattcccttttttgcggcattttg
ccttcctgtttttgctcacccagaaacgctggtgaaagtaaaagatgctgaagatcagttgggtgcacgag
tgggttacatcgaactggatctcaacagcggtaagatccttgagagttttcgccccgaagaacgttttcca
atgatgagcacttttaaagttctgctatgtggcgcggtattatcccgtattgacgccgggcaagagcaact
cggtcgccgcatacactattctcagaatgacttggttgagtactcaccagtcacagaaaagcatcttacgg
atggcatgacagtaagagaattatgcagtgctgccataaccatgagtgataacactgcggccaacttactt
ctgacaacgatcggaggaccgaaggagctaaccgcttttttgcacaacatgggggatcatgtaactcgcct
tgatcgttgggaaccggagctgaatgaagccataccaaacgacgagcgtgacaccacgatgcctgtagcaa
tggcaacaacgttgcgcaaactattaactggcgaactacttactctagcttcccggcaacaattaatagac
tggatggaggcggataaagttgcaggaccacttctgcgctcggcccttccggctggctggtttattgctga
taaatctggagccggtgagcgtgggtctcgcggtatcattgcagcactggggccagatggtaagccctccc
gtatcgtagttatctacacgacggggagtcaggcaactatggatgaacgaaatagacagatcgctgagata
ggtgcctcactgattaagcattggtaactgtcagaccaagtttactcatatatactttagattgatttaaa
acttcatttttaatttaaaaggatctaggtgaagatcctttttgataatctcatgaccaaaatcccttaac
gtgagttttcgttccactgagcgtcagaccccgtagaaaagatcaaaggatcttcttgagatcctttttt
ctgcgcgtaatctgctgcttgcaaacaaaaaaaccaccgctaccagcggtggtttgtttgccggatcaaga
gctaccaactctttttccgaaggtaactggcttcagcagagcgcagataccaaatactgttcttctagtgt
agccgtagttaggccaccacttcaagaactctgtagcaccgcctacatacctcgctctgctaatcctgtta
ccagtggctgctgccagtggcgataagtcgtgtcttaccgggttggactcaagacgatagttaccggataa
ggcgcagcggtcgggctgaacggggggttcgtgcacacagcccagcttggagcgaacgacctacaccgaac
tgagatacctacagcgtgagctatgagaaagcgccacgcttcccgaagggagaaaggcggacaggtatccg
gtaagcggcagggtcggaacaggagagcgcacgagggagcttccagggggaaacgcctggtatctttatag
tcctgtcgggtttcgccacctctgacttgagcgtcgatttttgtgatgctcgtcaggggggcggagcctat
ggaaaaacgccagcaacgcggcctttttacggttcctggccttttgctggccttttgctcacatgttcttt
cctgcgttatcccctgattctgtggataaccgtattaccgcctttgagtgagctgataccgctcgccgcag
c

### Equations

$$\begin{array}{c}\text{Normalized}\\ \text{Cas9 specificity}\\ \text{without Acr}\end{array} = \frac{\text{On/OT Cas9}}{\text{On/OT Cas9}} = 1 \tag{S1}$$

$$\begin{array}{c}\text{Normalized}\\ \text{Cas9 specificity}\\ \text{with Acr}\end{array} = \frac{\text{On/OT Acr–inhibited Cas9}}{\text{On/OT Cas9}} = 1.10 - 1.41 \tag{S2}$$

$$\begin{array}{c}\text{Increase in}\\ \text{Cas9 specificity}\\ \text{with Acr}\end{array} = \frac{\begin{array}{c}\text{Normalized}\\ \text{Cas9 specificity}\\ \text{with Acr}\end{array} - \begin{array}{c}\text{Normalized}\\ \text{Cas9 specificity}\\ \text{without Acr}\end{array}}{\begin{array}{c}\text{Normalized}\\ \text{Cas9 specificity}\\ \text{without Acr}\end{array}} \times 100\% = 10 - 41\% \tag{S3}$$
